# Spatiotemporal analysis of pericyte-induced dynamics in nascent angiogenic morphogenesis and heterogeneity using a microvessel-on-a-chip platform

**DOI:** 10.1101/2025.02.02.634898

**Authors:** Hedele Zeng, Takanori Sano, Jun-ichi Kawabe, Yukiko T. Matsunaga

## Abstract

The recent emergence of in vitro microvessel models has boosted research on angiogenesis and vascularization in tissue engineering, due to their superior control over in controlling culture conditions and enabling real-time observation compared to in vivo models. However, conventional two-dimensional (2D) observation and analysis fail to capture the heterogeneity of three-dimensional (3D) morphological dynamics. To overcome this issue, here, a novel morphological registration method for spatiotemporal quantification of angiogenic deformation dynamics by combining confocal microscopy of an engineered microvessel with computer vision techniques has been proposed in this paper. Using a coculture system of a microvessel and pericytes, the spatiotemporal measurements reveals: (i) distinct deformation patterns and growth/regression zonation on the parent vessel and angiogenic sprouts; (ii) spatiotemporal changes in pericyte localization and coverage; and (iii) the enhancing effect of pericyte-microvessel contact on local Notch signaling activation, the distribution of matrix metalloproteinase-1 (MMP-1), the heterogeneity of angiogenic dynamics, and morphological maturation. This pilot system offers a characterization of the comprehensive effects of cocultured cells during angiogenesis and enables the interactive fusion of multimodal data in future studies on vascular morphogenesis.

## 1. Introduction

Angiogenesis, the formation of new capillaries from pre-existing vessels, is a crucial factor in treating ischemic diseases^[1]^ and a key phenomenon in tumor growth and metastasis^[2]^. Numerous time-lapse studies have shown that elongation, stagnation, branching, fusion, and intussusception occur during the highly dynamic process of physiological angiogenesis^[3–5]^, while regression and pruning occur in pathological or conditioned angiogenesis^[6,7]^. However, existing high-resolution intravital microscopy systems are primarily designed for experimental animals, and species differences in angiogenesis pathways remain a challenge in conventional animal models^[8]^. Conventional noninvasive vascular imaging methods, such as computed tomography and dynamic magnetic resonance imaging, are mainly developed for observing large-scale angiogenesis phenomena^[9]^.

Recently, micro-physiological systems (MPSs) have garnered attention as potential in vitro experimental platforms for drug screening and developmental biology investigations. MPSs enable the recapitulation of tissue- and organ-level structures using multiple cell types, requiring less financial and time investment compared to traditional animal experiments. Among MPSs, models that mimic in vivo microvessel structures and functions provide researchers with a platform to easily characterize molecule transportation and vascular barrier function^[10,11]^ or observe angiogenic dynamics in real-time^[12,13]^. Due to the small thickness and high transparency of chip materials, such as glass and polydimethylsiloxane (PDMS), researchers can use confocal laser scanning microscopy^[14]^ or optical coherence tomography ^[15]^ to visualize, reconstruct, and retrace the inner vascular structure non-invasively at high resolution.

Nevertheless, although many in vitro microvessel models offer a three-dimensional (3D) culture environment and time-series 3D images, most angiogenesis studies are still primarily quantified using simple sprout counts or 2D parameters (e.g., sprout invasion distances and sprout diameter on maximum intensity projection (MIP) images). The variability and heterogeneity of angiogenesis profiles, influenced by differences in the surrounding environment, such as the three-layered morphogenesis in subretinal neovascularization, are easily overlooked^[16]^. The potential of these 3D culture models has not been fully explored, and there remains a need to develop analytical techniques for quantifying dynamic angiogenic behaviors without losing morphological information in space and time.

Three-dimensional registration is a process that transforms one 3D image (e.g., volumetric z-stack images) or geometry (e.g., triangular surface meshes or point cloud datasets) to align it with another related image or geometry. Normally, registration can be classified into rigid and non-rigid registration. Rigid registration allows translation, rotation, and scaling of an object to register, while nonrigid registration also accepts flexible transformation^[17]^. This irregular transformation results in high morphological similarity between the registered and fixed datasets, enabling a distance-based point-to-point pairing between them. Some studies have demonstrated the effectiveness of this registration method by comparing point cloud datasets generated from time-series 3D CT-scanned thoracic aortic aneurysm images^[18]^ or 3D point clouds obtained during plant growth^[19]^. However, conventional free-form deformation methods used in biomedical image studies are mainly developed to map isometric or near-isometric shapes^[20]^ and predict smooth and continuous deformation fields^[21]^. This characteristic makes it difficult to simultaneously handle large-scale non-isometric angiogenic changes within large time intervals. Skeletonization-based registration still needs other models, such as hidden Markov models or Bayesian networks, to predict the relationships between nodes. Specifically for angiogenesis measurement, skeletonization only extracts the central line of angiogenic sprouts, resulting in the loss of important surface features, such as local smoothness and small protrusions on the parent vessel surface^[22]^.

Here, a novel morphology-adaptive Gaussian radial basis function-based iterative non-rigid registration (MAGIN) method was applied to quantify morphological changes on a capillary-resident pericyte cocultured in vitro microvessel model, which is designed to recapitulate the angiogenesis process following regenerative stem cell implantation. Briefly, 3D microvessel surface meshes reconstructed from confocal z-stack images were registered in the time series to quantify the deformation of the microvessel surface at each position. Concurrently, we used Fourier transform and filtering to divide the parent vessel and sprout areas of the microvessel, providing spatial information to guide the registration grid size. The location of pericytes was determined by their overlap with the microvessel structure. By integrating the deformation of endothelial cells (ECs) with the relative position of pericytes to ECs, the diverse effects of pericyte coverage on vascular morphology were deduced. Analysis results showed that: 1) The sprout area had a greater deformation distance and a more distinct growth/regression zonation along the radial direction than the parent vessel. 2) Pericytes selectively adhered to the sprout area and were less likely to detach. 3) Pericyte coverage promoted sprout differentiation and accelerated the maturation of the entire sprout. To our knowledge, while these angiogenic dynamics have been observed or qualitatively described in numerous previous studies, they have not been quantitatively measured from the perspective of vascular surface dynamics. We believe this novel non-rigid registration method, derived from computer vision and specifically shape matching techniques, offers a new perspective for future in vitro angiogenesis studies and can contribute to therapeutic angiogenesis research.

## 2. Results

### 2.1. Recapitulation of the angiogenic morphogenesis process using an in vitro microvessel model

We fabricated the microvessel chip using endothelial cells (ECs) and pericytes mixed in a collagen gel, as previously described (Figure 1a)^[14]^. Our previous study demonstrated that this peripheral tissue capillary-resident EphA7+ pericyte rescues blood flow in vivo^[23]^, enhances sprouting angiogenesis^[14]^, and maintains vascular barrier function in vitro^[24]^. Microscopic images revealed significant vessel expansion and vessel-directed pericyte migration from day 1 (1 day after cell seeding) to day 5 (Figure 1b). By day 5, we observed angiogenic sprouts (white arrows) and loop-like structures formed by the fusion of multiple sprouts (white arrowhead).

**Figure 1.**
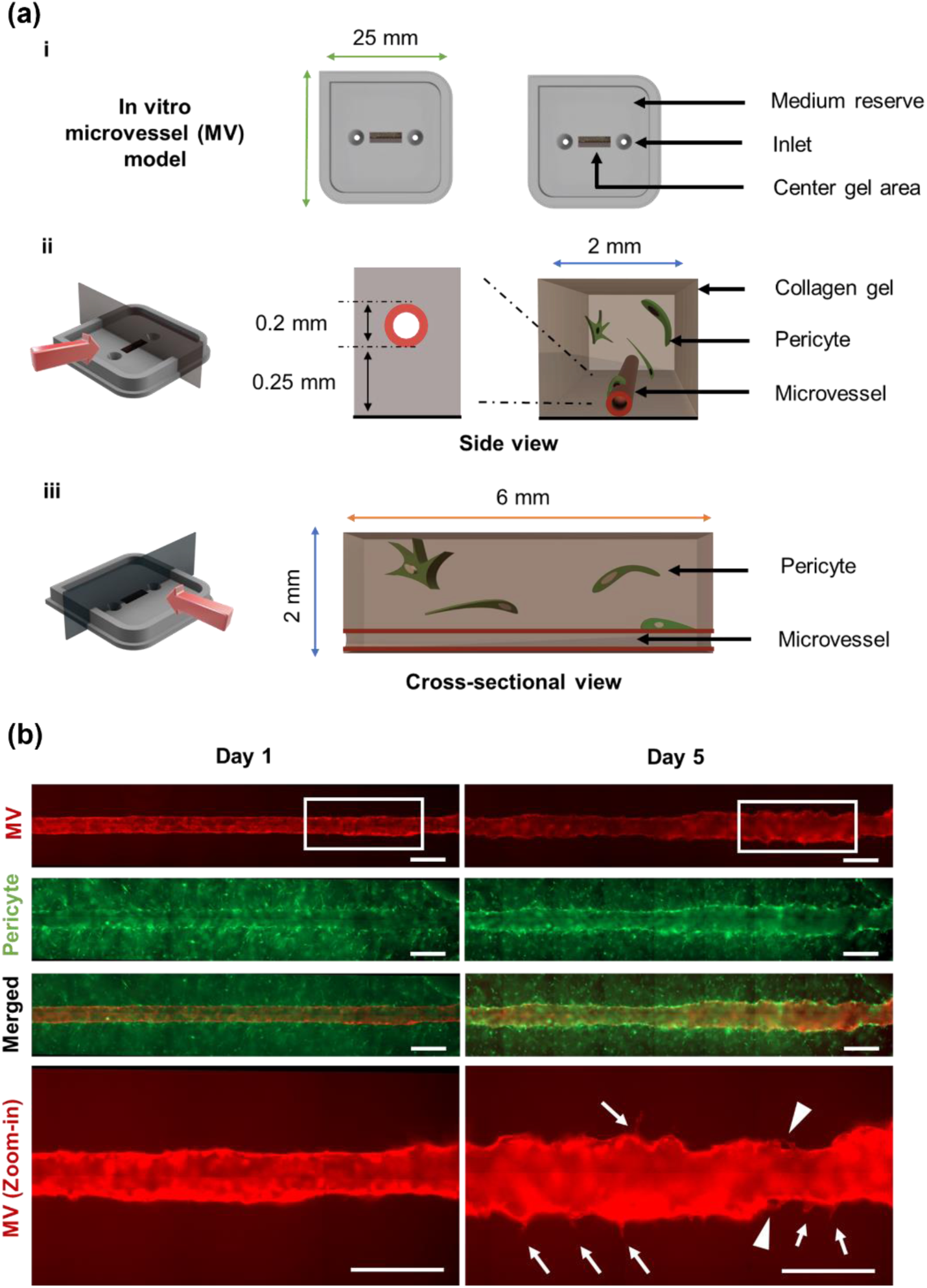
Construction of the in vitro microvessel model and angiogenesis observation. **a)** Schematic illustration of the in vitro microvessel chip (i), the side view (ii) and the cross-sectional view (iii) of the center gel area. **b)** Time-lapse fluorescent images of microvessel and surrounding pericytes. A representative angiogenic area, highlighted by the white rectangle, is shown at higher magnification below. In the day 5 image, white arrows and white arrowheads indicate sprouts and fused ring-like structures, respectively. Scale bar: 400 μm.

### 2.2. Morphology-adaptive Gaussian radial basis function-based iterative nonrigid registration enables time-series mesh tracking and high-resolution angiogenic dynamic measurement

We used confocal laser scanning microscope (CLSM) and acquired time-series z-stack 3D images of the in vitro microvessel model for quantitative angiogenesis analysis. The 3D images were then processed by IMARIS (Bitplane, Zurich, Switzerland) to reconstruct the surface mesh, for the subsequent discrete vertex coordinate extraction (Figure 2a-ii), external surface extraction (Figure S1), parent vessel-sprout classification and mesh pre-alignment. Local deformations were calculated on control points and applied to source vertices to align the two objects morphologically. The registered point positions indicate the correspondence between the registered mesh vertices and the broadline. This allows us to determine the correspondence, calculate deformation vectors, and estimate local morphological changes in the microvessel.

**Figure 2.**
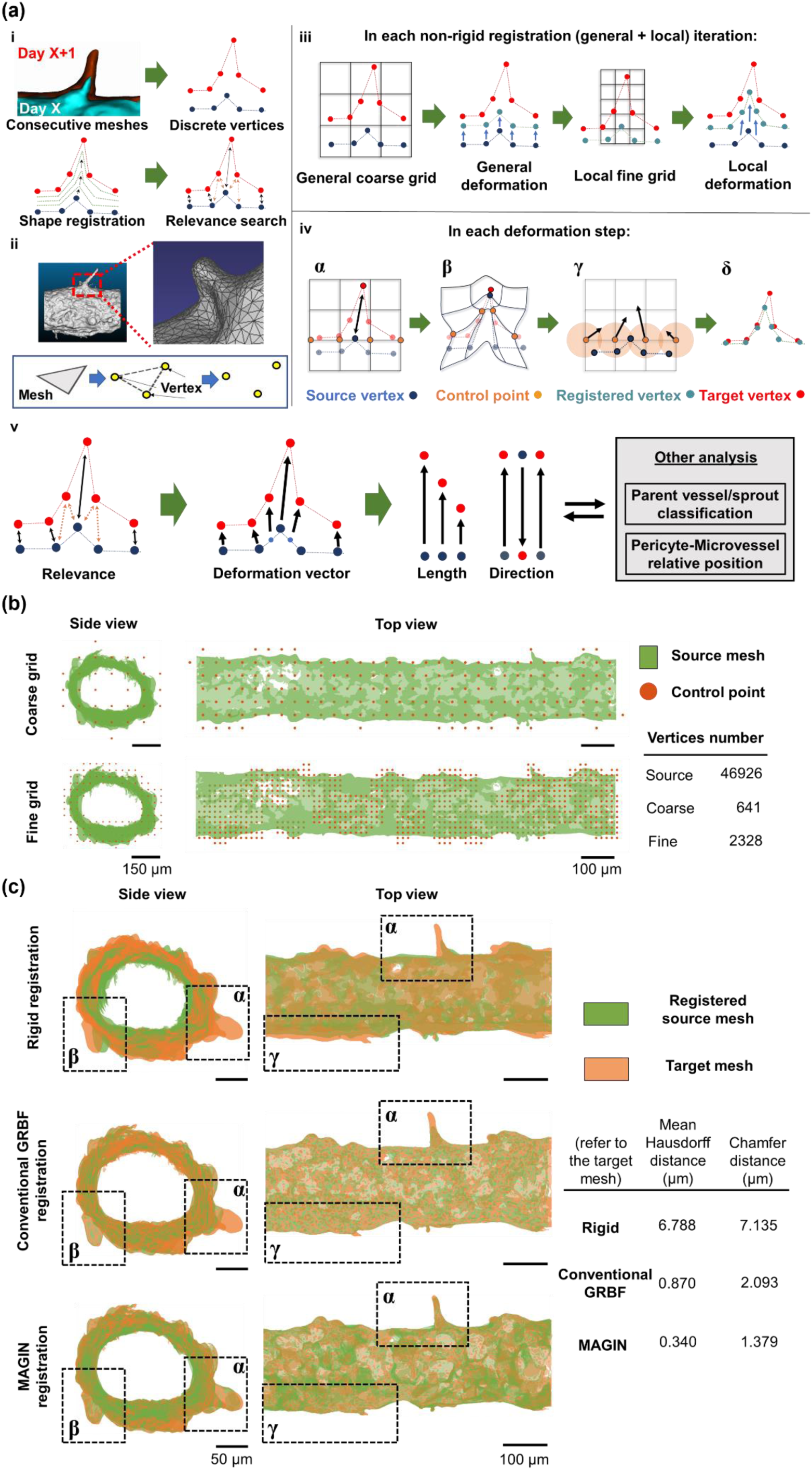
Non-rigid deformation for time-series mesh registration. **a)** Schematic illustration of vertex data extraction, shape registration, and vertex correspondence search (i). Vertex coordinates were extracted from the surface mesh (ii). In the shape registration procedure, each iteration contains a coarse grid-controlled general deformation and a fine grid-controlled local deformation, particularly in sprout areas (iii). Each deformation step contains α. virtual grid creation, control point selection, and morphological constraint definition; β. control point deformation vector calculation; γ. deformation interpolation; and δ. mesh splitting and local adjustment (iv). Calculation of deformation distance and direction, followed by analysis with other information (v). **b)** Comparison between the coarse grid for general deformation and the fine grid for local deformation. **c)** Target data and registered data processed using only rigid registration (top), conventional GBRF registration (middle), and the proposed registration method (MAGIN) (bottom). A representative surface mesh set from day 3 and day 4 was presented as “source mesh” and “target mesh.” Representative areas are highlighted by the dotted line boxes. Units: μm. Mean Hausdorff and Chamfer distances with respect to the target mesh after registration are shown.

To decrease the execution time per iteration, the mesh deformation was defined by a small number of control points, derived from the nodes of a virtual grid within the 3D bounding box of the mesh. Parent vessel-sprout segmentation was initially performed using a Fourier transform-based method we previously proposed^[14]^. In each iteration of the non-rigid registration procedure, there were two deformation processes: a coarse grid was used for global deformation, and a fine grid was used for local, accurate deformation in the sprouting area (Figure 2a-iii). This morphology-adaptive registration strategy was developed to address large-scale non-isometric deformation during angiogenesis and to balance deformation accuracy with computational cost. In each deformation step, control points were selected from the virtual grid nodes around the source vertices. The weights from control points to source vertices were calculated using a Gaussian radial basis function (GRBF) and the Euclidean distance between them. Concurrently, morphological constraints between source and target vertices were defined (Figure 2a-iv-a). Based on these weights, a set of deformation vectors was calculated for each control point after Tikhonov regularization^[25]^ to satisfy the constraints while minimizing the overall residual error^[26]^ (Figure 2a-iv-b). The deformation vectors at each control point were then interpolated to all source vertices, creating a smooth elastic deformation field^[27]^ (Figure 2a-iv-c). Local deformation in the sprout area tends to be large-scale. Triangle meshes with a relatively large aspect ratio and square were split at the end of each iteration to prevent over-stretching of individual triangles and to provide new vertices for further extension (Figure 2a-iv-d). Grid resolution and the GRBF shape parameter were iteratively increased in an annealing process. The local position was adjusted based on the normal direction and neighborhood of each source vertex at the end of each iteration^[28]^. Using the correspondence and original positions of the registered mesh vertices, we calculated the deformation vector for each vertex on the target mesh and interpolate the deformation (Movie S1). The length and direction of deformation vectors were further integrated with other geometrical information to quantify the heterogeneity of the microvessel morphology change during angiogenesis (Figure 2a-v).

Here, we selected a representative mesh dataset to demonstrate the grid refinement process and compare registration methods. The grid was constructed with a higher resolution only in the sprout area, while the total number of nodes remained significantly smaller than the number of source vertices (Figure 2b). The comparison of registration methods in Figure 2c, Figure S3 and Figure S4 showed that nonrigid registration, flexible GRBF registration especially, achieved better surface fitting than rigid registration alone in dealing with curved structure and irregular deformation The overall deformation defined by rigid registration was incompatible with the irregular nature of angiogenic morphogenesis. Furthermore, we found that the conventional GRBF method was suitable for large-area deformations, as highlighted in regions β and γ in Figure 2c, but had limitations in sprout areas with long-distance deformations, as highlighted in region α in Figure 2c. The coarse grid used in conventional GRBF registration made the control points less sensitive to large deformation components, ultimately resulting in a homogeneous deformation field on the source vertices. In contrast, the MAGIN method generated a dense set of control points around the sprout areas. The high density and local deformation characteristics allowed for a higher GRBF shape parameter setting, enabling the generation of a deformation field with sufficient heterogeneity at the sprout vertices. The mean Hausdorff distance and Chamfer distance between the registered meshes and the target mesh also quantitatively demonstrated the superiority of the MAGIN method in addressing angiogenic deformation. Improved surface fitting and good morphology preservation made distance-based correspondence searching feasible and reliable.

### 2.3 Local dynamics type zonation on angiogenic sprouts: growth tip area and regression at the root area

Local regression and growth status were defined based on correspondence in time-series data, and their spatial distributions were analyzed. We assumed the normal vector direction as the growth direction for each vertex. The angle between a vertex’s normal and its deformation vector was calculated to measure local growth and regression. Vertices with an angle greater than 90° were considered to exhibit growth with respect to their local surface mesh area; conversely, vertices with an angle less than 90° were considered to be in a state of regression (Figure 3a). Distribution histograms show that the majority of vertices for both parent vessel and sprout vertex groups were in the growth zone (0° < 𝜃 < 90°), indicating that the sample microvessel generally exhibited growth during the observation period. When focusing on each zone, we found a greater number of parent vessel vertices in the regression zone (90° < 𝜃 < 180°), while a larger proportion of sprout vertices were in the growth zone (0° < 𝜃 < 90°) (Figure 3b). The colorized growth state distribution map of the representative data also exhibited a similar trend. The growth area (in green) predominated over the regression area (in blue) in both the parent vessel and sprout (Figure 3c). We considered the larger proportion of growth zone in sprout area contributed to a higher mean deformation distance than that of the parent vessel (Figure S5). Intriguingly, the growth and regression areas appear randomly distributed across the parent vessel surface mesh, whereas in the sprout area, the regression vertices were predominantly located at the root area of the sprouts (Figure 3c-ii).

**Figure 3.**
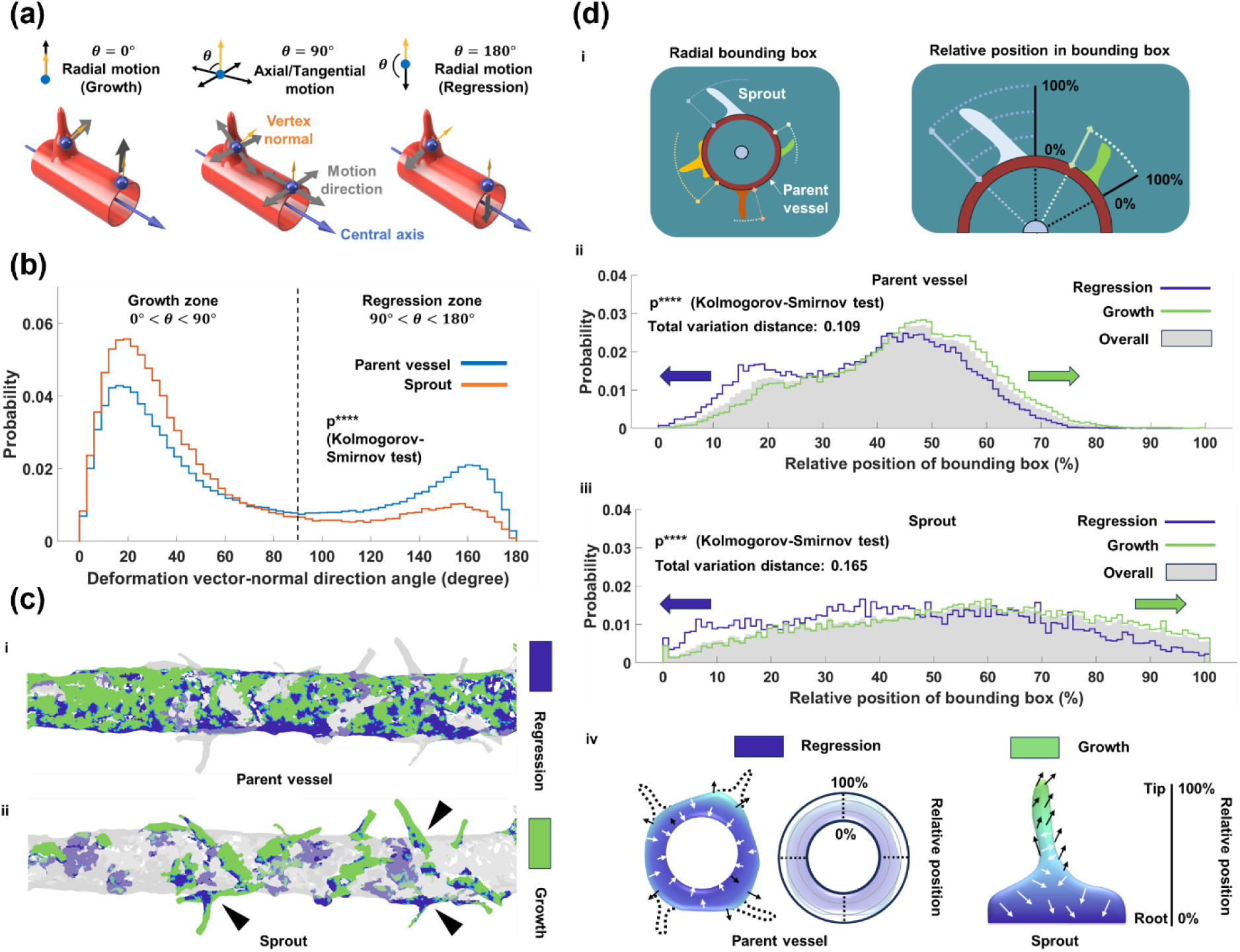
Growth/regression state classification and comparison between parent vessel and sprout. **a)** Schematic illustration of vertex local motion types, defining “growth” and “regression.” **b)** Probability distribution histogram of the angle between the deformation vector and the vertex normal direction. All surface mesh vertices from seven samples were combined to draw the histogram. **c)** Growth/regression state map of a representative surface mesh: parent vessel area (i) and sprout area (ii). Three sprouts with regressive regions at their bases are highlighted by black arrowheads. A representative surface mesh from day 5 and its corresponding deformation data (from day 4 to day 5) were presented. **d)** Definition of the bounding box along the vessel’s radial direction and the relative position within the bounding box (i). Probability distribution histogram of the relative position within the bounding box. Histograms (ii-iii) were generated using a general distribution for all vertices and specific distributions for growth and regression vertices on both the parent vessel and sprout. Schematic illustration of the spatial zonation of the growth/regression state in relation to the bounding box (iv). p**** means p ≤ 0.0001.

We further examined the spatial distribution features of the growth/regression state (Figure 3d). We extracted the principal component analysis (PCA) first axis of inner surface of the source vessel mesh and defined this as the vessel central axis. We then obtained vectors originating from the central axis, pointing to each sprout vertex, and perpendicular to the axis. The minimum and maximum distances from each sprout to the central line were recorded. We defined these limits as the boundaries of the bounding box in the radial direction (Figure 3d-i).

The growth histogram closely resembled the general distribution shape in the parent vessel and sprout results, consistent with the dominance of the growth state in Figure 3B. Moreover, both datasets depict a separation between the regression and growth histograms, as indicated by the relative position within the bounding box (Figure 3d-ii&iii). We used total variation distance to quantify the difference between the two probability distributions (regression and growth). Numerical results show that the largest absolute difference between the probabilities was larger in the sprout area (0.165) than in the parent vessel (0.109), indicating that both the parent vessel and sprout had regression and growth, with this phenomenon being more pronounced in sprouts (Figure 3d-iv).

### 2.5 Pericyte selectively migrates to sprout, increases sprout heterogeneity and promotes sprout morphological maturation via Notch and MMP-1 regulation

Our previous studies have shown that peripheral tissue capillary-resident EphA7+ pericytes are superior in blood flow rescue^[23]^, vascular barrier function enhancement^[24]^, and angiogenic sprout morphological maturation^[14]^. Here, we extracted the relative positions of pericytes and microvessel from the time-course dataset and integrated this information with i) parent vessel-sprout classification labels, ii) vertex deformation information, and iii) temporal changes in the morphology of specific sprouts. This was done to reveal the spatiotemporal dynamics of pericyte residence and pericyte-microvessel crosstalk (Figure 4).

**Figure 4.**
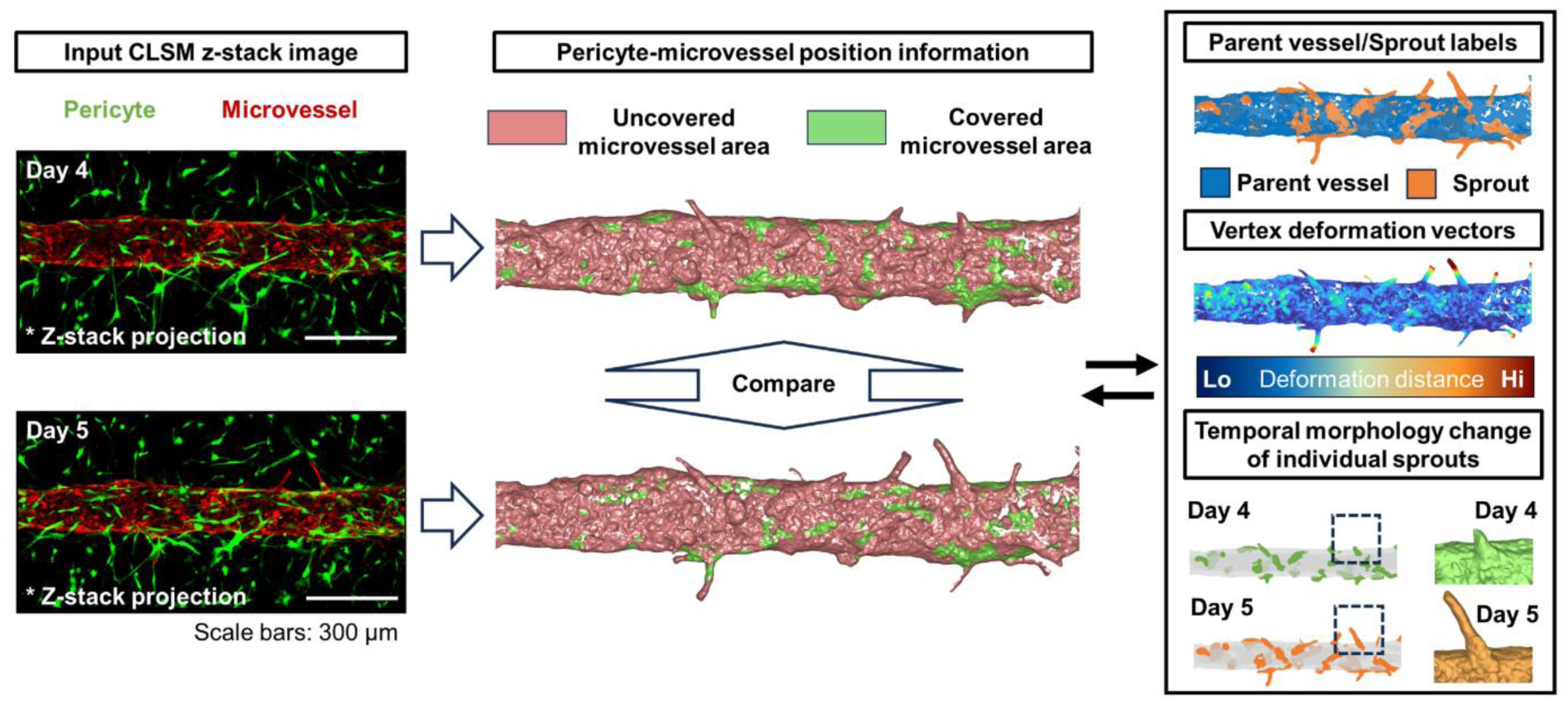
Schematic illustration of pericyte-microvessel relative position information extraction and other information types combined in the following analysis.

We first surveyed the temporal change in the “pericyte-covered vertex” ratio among all vertices, especially in the sprout area (Figure 5). microvessel vertices encapsulated within the pericyte surface mesh were labeled as “pericyte-covered vertices”; otherwise, they were labeled as “pericyte-uncovered vertices.” Pericyte coverage data for each sample microvessel were continuously tracked from day 3 to day 5. Results show a slight general increase in coverage rate, with averages starting from 8.03% (day 3) to 9.90% (day 4) and 9.98% (day 5) (Figure 5a). The sprout vertex ratio among all pericyte-covered vertices was calculated to assess potential bias in pericyte migration. Results show a significant increase in the average sprout vertex ratio, from 3.80% on day 3 to 18.4% on day 4, followed by a smaller increase to 19.7% on day 5 (Figure 5b). For the pericyte-covered vertices, we also examined the preservation rate of the coverage state. If a pericyte-covered vertex on the previous day and its corresponding vertex on the subsequent day were also classified as pericyte-covered, we counted the state of this vertex as “preserved.” We found that in both time periods, day 3 to day 4 and day 4 to day 5, the mean preservation ratio of the sprout area (day 3 to day 4: 33.7% and day 4 to day 5: 43.6%) was higher than the general level (day 3 to day 4: 31.6% and day 4 to day 5: 39.6%), suggesting that pericytes are more likely to remain in the sprout area (Figure 5c). Based on this evidence, we believe that there is a sprout bias in pericyte migration and settlement (Figure 5d).

**Figure 5.**
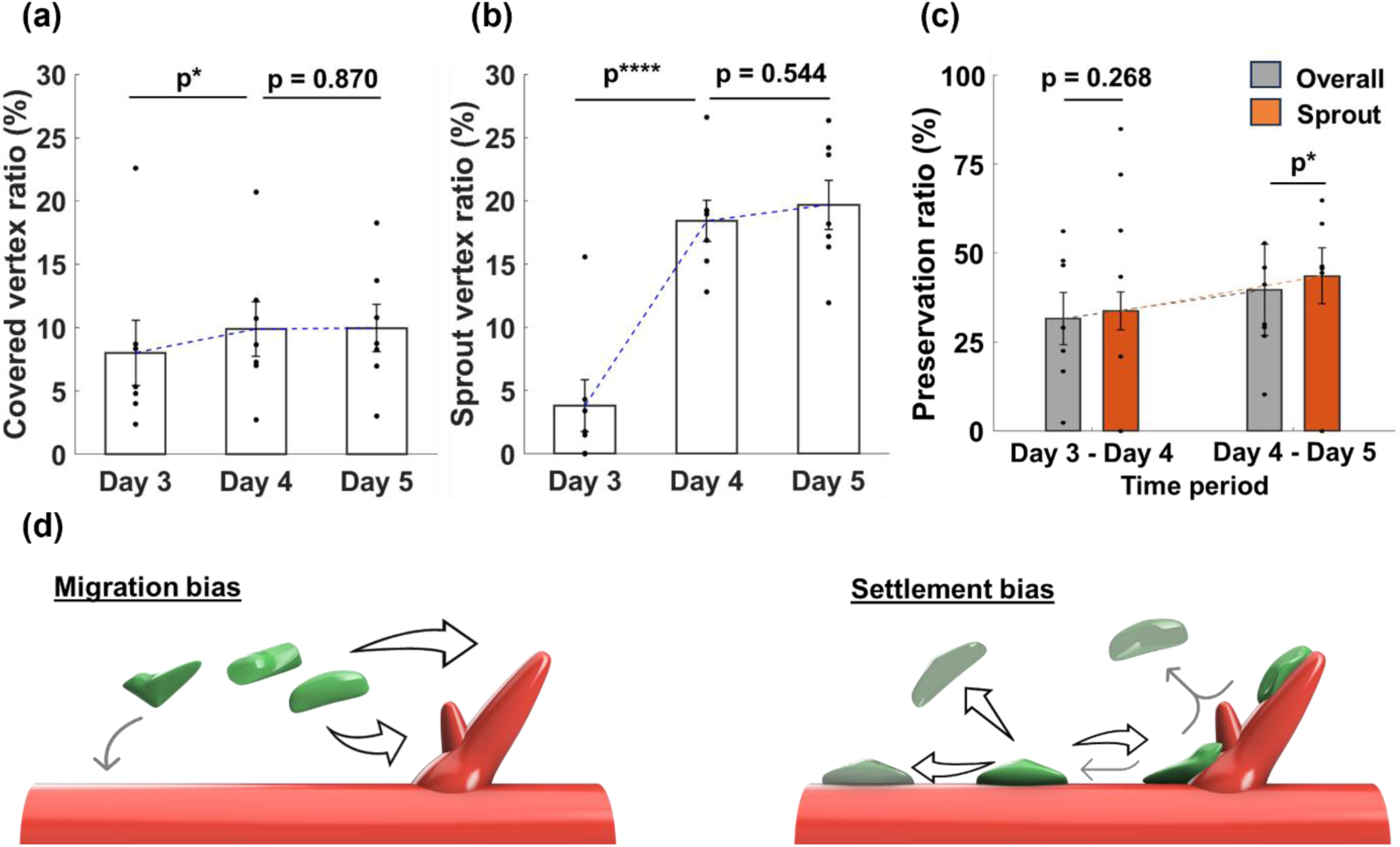
Quantitative analysis of pericyte coverage state temporal change. **a)** Temporal change of the “pericyte-covered vertex” ratio among all the surface mesh vertices. **b)** Temporal change of the sprout vertex ratio among all the “pericyte-covered vertices” in each day. **c)** Temporal change of the preservation ratio of the “pericyte-covered” state in different time periods and area types. If both a vertex and its corresponding vertex on the subsequent day are classified as “pericyte-covered vertices,” we define this as “preserved.” If a vertex is classified as “pericyte-covered vertex” and its corresponding vertex on the subsequent day is classified as “pericyte-uncovered vertex,” we define this as “unpreserved.” Preservation ratio was calculated as the number of “preserved” vertex divided by the total number of pericyte-covered vertex on the previous day. Data are shown as mean ± standard deviation. **d)** Schematic illustration of the sprout-bias during pericyte migration and the coverage preservation preference on sprout area. p-values were from paired permutation tests respectively. p*: p ≤ 0.05, p****: p ≤ 0.0001

We observed a clear difference between pericyte-covered and pericyte-uncovered vertices after integrating pericyte coverage state with the deformation vector (Figure 6). The deformation distances of pericyte-covered vertices were more concentrated in the small deformation zone (0-10 μm), while those of pericyte-uncovered vertices had a more even distribution and a larger share in the larger deformation zone (10–40 μm) (Figure 6a). Meanwhile, we used the same “growth/regression” deformation state measurement as in section 2.4. The distribution plot of the deformation vector-vertex normal angle shows that the majority of both pericyte-covered and pericyte-uncovered vertices were concentrated in the growth zone (0° < *θ* < 90°), with pericyte-covered vertices showing a slight advantage, although the two probability lines overlapped in the central part. Conversely, pericyte-uncovered vertices were predominant in the right regression zone (90° < *θ* < 180°) (Figure 6b). We further integrated the parent vessel-sprout classification information into the analysis. Results show both pericyte-covered and pericyte-uncovered sprout vertices exhibit the growth/regression zonation observed in section 2.4. However, the mean relative position of pericyte-covered sprout vertices (48.7%) is closer to the sprout root than that of pericyte-uncovered sprout vertices (52.3%). The distribution of regression vertex position is also more concentrated and differs from the growth group under pericyte-covered conditions (maximum absolute difference between empirical cumulative distribution functions (CDFs): 0.387, total variation distance (TVD): 0.462) compared to pericyte-uncovered conditions (maximum absolute difference between empirical CDFs: 0.149, total variation distance (TVD): 0.164) (Figure 6c & 6d). In contrast, the difference between pericyte-covered and pericyte-uncovered parent vessel vertices was unclear (Figure S6). To verify the local dynamic zonation with biological evidence and quantify the effects of pericytes, we visualized the Notch signaling activation marker, Notch-1 intracellular domain (N1ICD), and the vascular regression-related marker, matrix metalloproteinase-1 (MMP-1), on sprouts. Overall, pericyte-EC coculture chips had a higher EC N1ICD and MMP-1 expression level than that of EC monoculture chips (Figure S7). Spatially, images showed that pericyte-EC coculture chips had heterogeneous N1ICD and MMP-1 expression levels along the sprout, with high expression areas concentrated in the root area. In contrast, N1ICD and MMP-1 levels on EC monoculture chip sprouts were homogeneously distributed (Figure 6e & 6f). Fluorescence intensity along the sprout boundary was measured from root to tip and normalized across samples. Quantitative analysis revealed a steep decrease in N1ICD and MMP-1 expression levels from root to tip on coculture sprouts, and a uniform distribution with a slight decrease toward the tip on monoculture sprouts, consistent with the pattern observed in the representative images. Notably, MMP-1 showed a slight up-regulation at the tip (Figure 6g & 6h). Based on this evidence, we believe that pericyte-EC cell-cell contact contributes to the heterogeneity of N1ICD and MMP-1 expression level, enhances the contrast between root and tip ECs, regulates neovessel morphology, and promotes elongation (Figure 6i).

**Figure 6.**
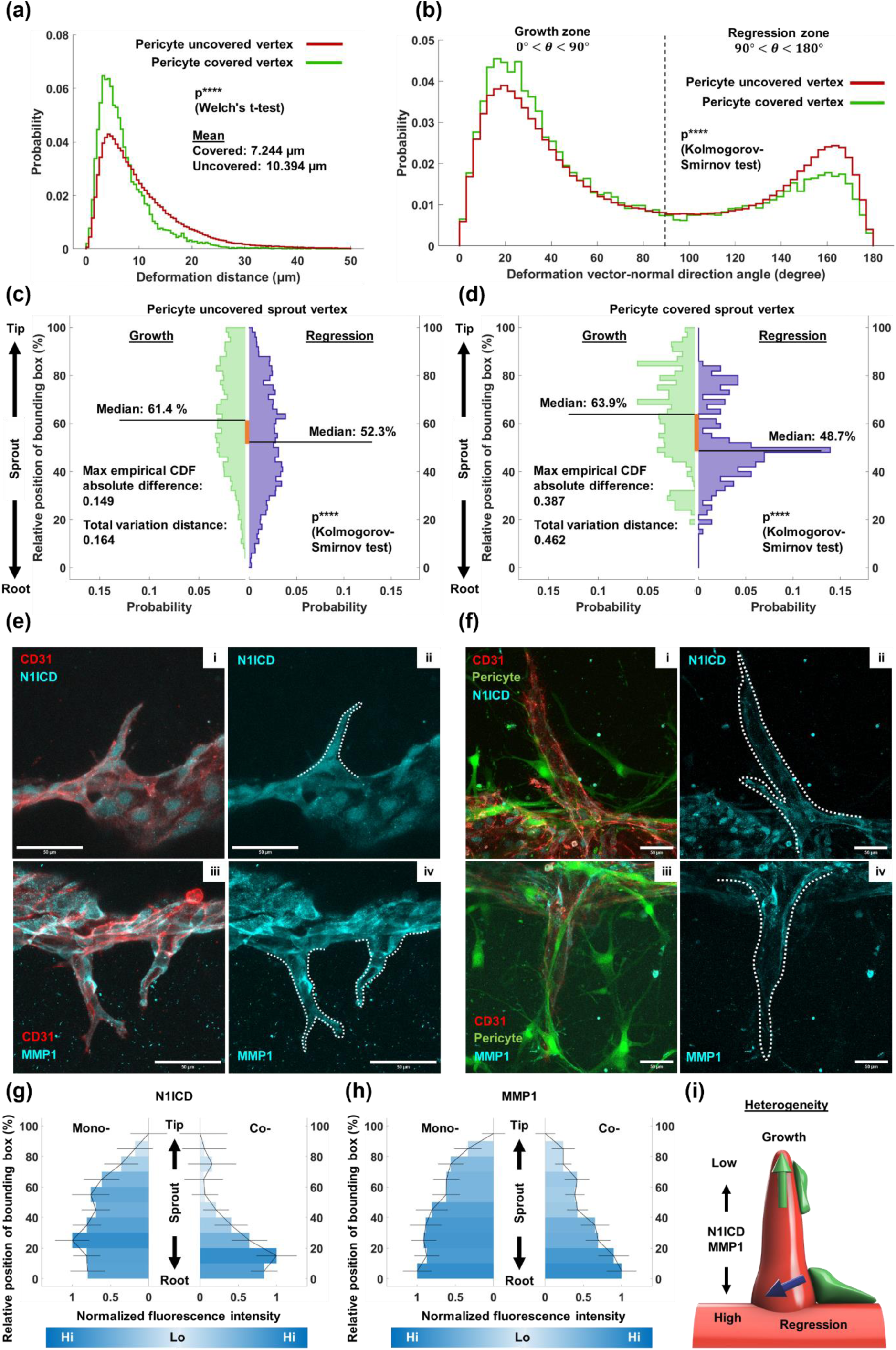
Comprehensive analysis of pericyte influence on angiogenic dynamics. Probability distribution histogram of **a)** the microvessel vertex deformation distance and **b)** the angle between the microvessel vertex deformation vector and the vertex normal vector. All the surface mesh vertices from seven samples were combined to draw the histogram. Only if a vertex meets two requirements, α) it is labeled as “pericyte-covered vertex”; β) its corresponding vertex on the subsequent day is also labeled as “pericyte-covered vertex,” was it classified as the “pericyte-covered vertex” in this measurement. The same applies to “pericyte-uncovered vertices.” Probability distribution histogram of relative position for **c)** pericyte-covered and **d)** pericyte-uncovered sprout vertices. Growth and regression vertices were separated to draw the histogram. Immunofluorescent images of **e)** sprouts from EC monoculture samples and **f)** sprouts from pericyte-EC coculture samples. Sprout boundaries were highlighted by the white dotted line. Spatial distribution of normalized fluorescence intensity of **g)** N1ICD and **h)** MMP1 along the sprout. The sprout root was set as relative position 0 and the tip was set as 100. Intensity was averaged among all samples, the highest value along the sprout was set as normalized fluorescence intensity 1, and the lowest value was set at normalized fluorescence intensity 0. Error bar: standard error. **i)** Schematic illustration of pericyte-EC contact influence on angiogenic morphogenesis. Scale bar: 50 μm.

To quantitatively measure the effects of pericyte coverage on sprout morphology, we paired the sprouts in the time-series data, selected sprouts that did not undergo regression or fusion during the observation period, examined their pericyte coverage state, and traced their temporal Gaussian curvature standard deviation (STD) and changes in sprout length (Figure 7).

**Figure 7.**
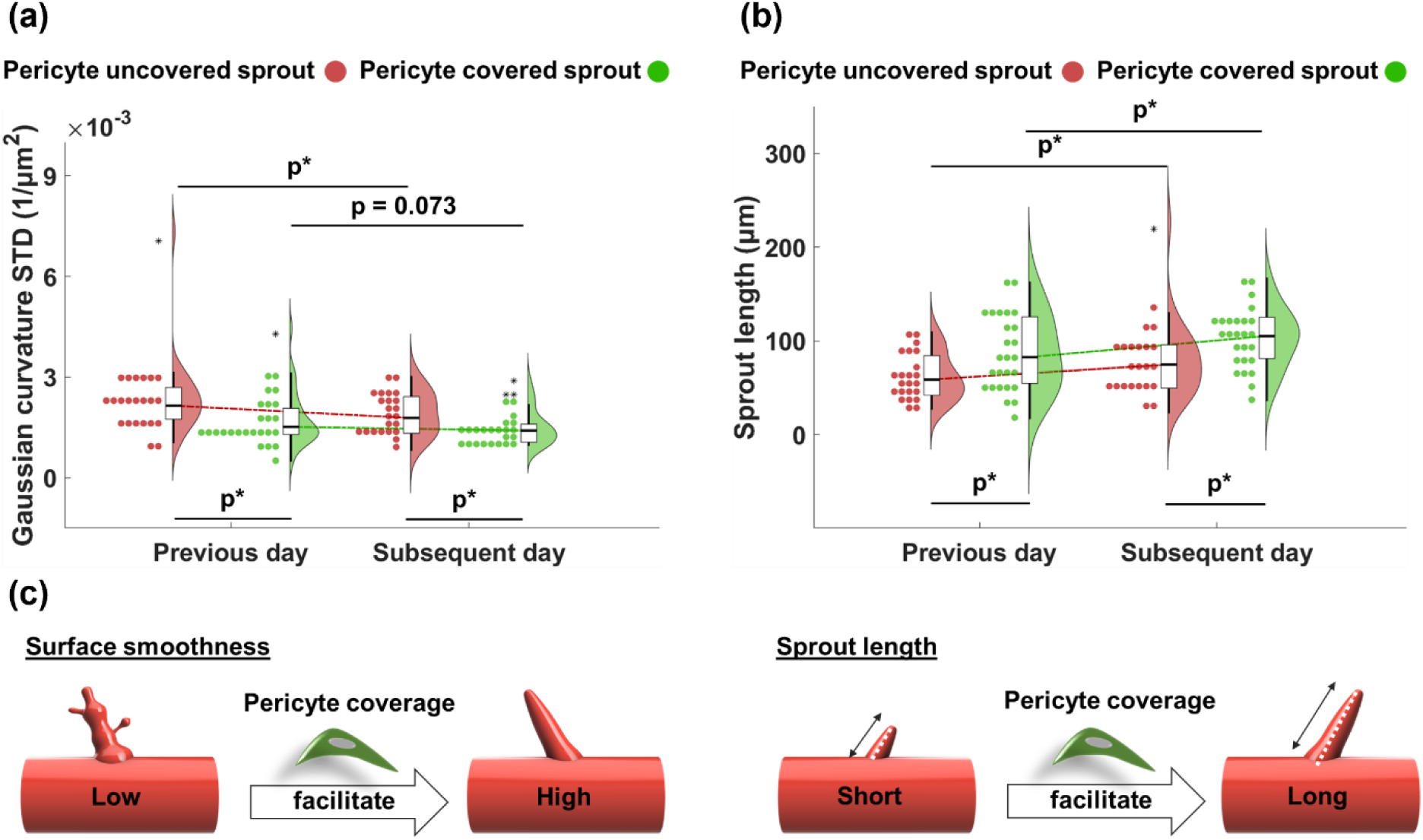
Quantitative analysis of pericyte coverage effects on individual sprouts morphology. Temporal change of **a)** sprout mesh Gaussian curvature standard deviation (STD) and **b)** sprout length. Only if both a sprout and its corresponding sprout on the other day contain “pericyte-covered vertices,” was it classified as the “pericyte-covered sprout.” The same applies to “pericyte-uncovered sprouts.” Data are represented as box plots to show medians. p-values were from paired (between different days) and unpaired (between pericyte-covered and -uncovered groups) permutation tests respectively. p*: p ≤ 0.05. **c)** Schematic illustration of the pericyte coverage effects on sprout smoothing and elongation.

Gaussian curvature is a widely used local concavity parameter and has been shown to have biological significance^[29]^. Here, we used the Gaussian curvature STD as a geometrical index of overall surface smoothness. A high Gaussian curvature STD indicates intense curvature changes and a rough surface, vice versa. Results show that there was a general decrease in the Gaussian curvature STD over time, while the medians of pericyte-uncovered sprout Gaussian curvature STD (previous day: 2.15 × 10^−3^, subsequent day: 1.78 × 10^−3^, unit: μm^−2^) remained larger than that of pericyte-covered sprouts (previous day: 1.52 × 10^−3^, subsequent day: 1.41 × 10^−3^, unit: μm^−2^), indicating that pericyte coverage contributes to the sprout smoothing process (Figure 7a).

Here, we defined the projection of the surface mesh onto its PCA first axis as the sprout length, since sprout growth in 3D space is stochastic and difficult to measure from only a single direction (radial, axial, or tangential). Similarly, we compared the temporal changes in length between pericyte-uncovered and pericyte-covered sprouts. Results show that both groups achieved sprout elongation over time, and the median of pericyte-covered sprout lengths (previous day: 82.6 μm subsequent day: 105 μm) remained larger than that of pericyte-uncovered sprouts (previous day: 58.7 μm subsequent day: 74.6 μm). The difference in the median values (pericyte-covered sprout: 22.5 μm; pericyte-uncovered sprout:16.0 μm) also suggested that pericyte coverage promotes the elongation of the sprout (Figure 7b). The morphological comparisons between pericyte-uncovered and -covered sprouts demonstrated that pericyte-microvessel contact is significant for sprout surface smoothing and angiogenic sprout elongation (Figure 7c). This effect may not be replicated or compensated for by other indirect, long-distance cell–cell interactions.

## 3. Discussion

In this study, we constructed a 3D pericyte-microvessel coculture system, proposed a novel morphology-adaptive Gaussian radial basis function-based iterative non-rigid registration (MAGIN) method for in vitro angiogenesis investigation, and quantitatively measured the influence of pericyte coverage on microvessel local dynamics and the nascent sprout maturation process. Numerical measurements of time-series 3D mesh provide new perspectives on angiogenesis study. Pericyte direct contact induced heterogenization and sprout root area regression, which were first discovered in this study, unraveling a part of the comprehensive morphogenesis process and verifying the accuracy of the MAGIN method (Figure 8).

**Figure 8.**
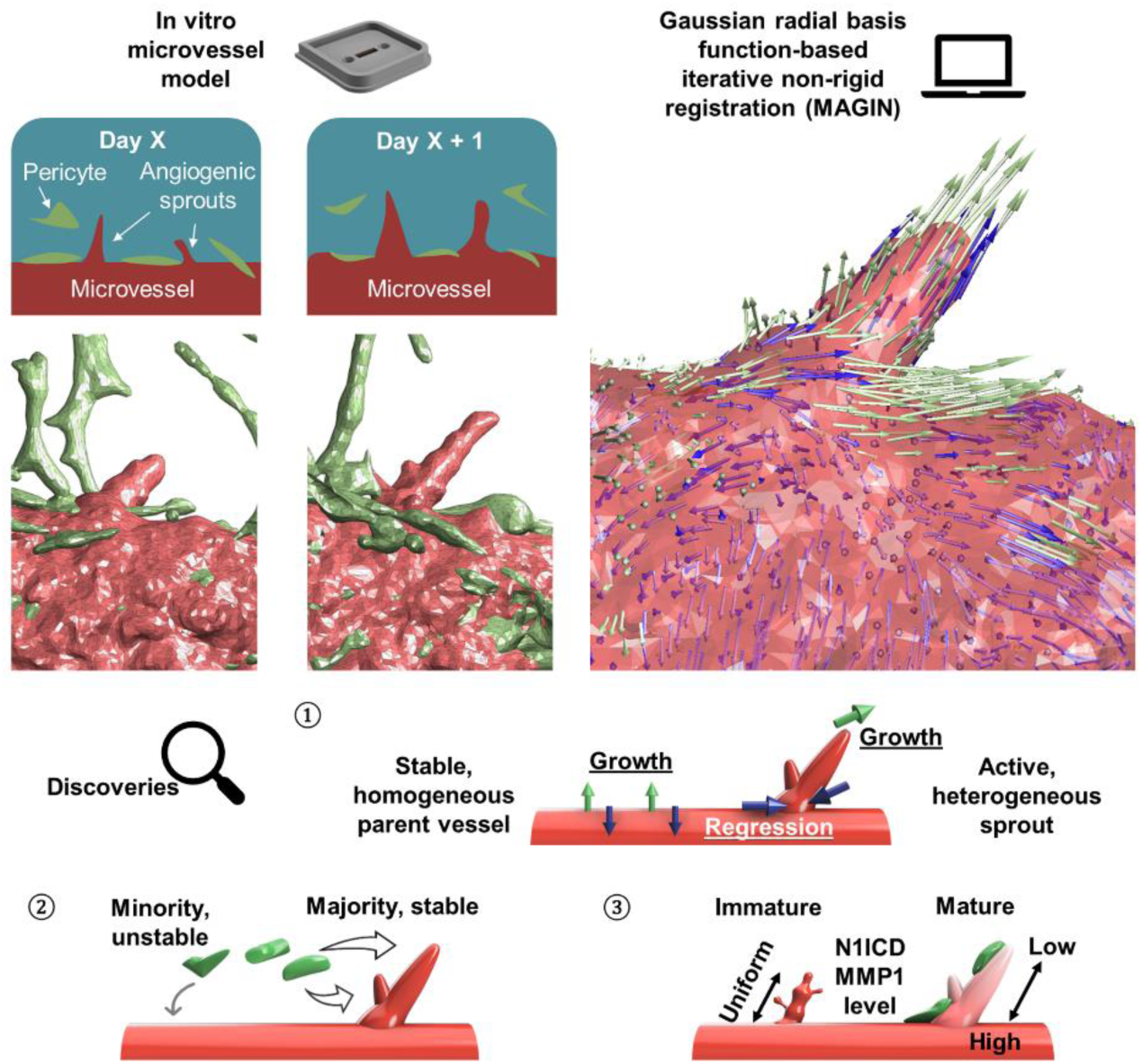
Schematic illustration of the angiogenic dynamics and pericyte-microvessel interaction investigation through non-rigid surface mesh registration.

The in vitro angiogenesis model, initially developed for investigating microvascular lumen formation and tumor angiogenesis factors^[30]^, is currently drawing attention from the tissue engineering field. This is because the physical cues and chemical factors screened by this model can contribute to the development of perfusable, ordered, and dense vascular networks in engineered tissues^[31]^. To meet this need, a quantitative method capable of analyzing spatiotemporal dynamics is required. Discussions and conventional 2D measurements, such as invasion distance^[22]^ and sprout branch number^[32]^, typically treat the entire sprout as a single object, providing only a general assessment of angiogenesis. In contrast, the deformation of a 3D mesh can be decomposed into the deformation of numerous discrete vertices. In this context, a 1-mm length region of interest (ROI) in a vessel mesh can contain approximately 80,000 vertices, with triangle mesh line lengths of approximately 5 μm.

Interestingly, if we consider this as a computer graphics shape matching problem, the pattern of angiogenic deformation is distinct from some benchmark models. It differs significantly from the alignment of different facial expressions^[33]^ or various human/animal gestures^[34,35]^. These are considered near-isometric problems, involving large deformations but preserving the features of specific landmark areas (e.g., nose, mouth, hands). In contrast, angiogenesis involves sprouts undergoing large-scale growth or regression and changing direction between time points. While emerging nonisometric shape mapping methods may address the issue of sprout growth/regression^[36]^, their applicability to specific situations, such as the emergence of new sprouts or the smoothing of small swelling points, requires further investigation. Some studies have employed free-form deformation based 3D elastic registration, however, the deformation patterns of the objects in these studies, such as those in tumor^[37]^ and bone tissue^[38]^, are still not as non-isometric as the angiogenesis pattern. Skeletonization and subsequent feature extraction remain the primary approaches in high-dimensional angiogenesis^[32]^ and vessel remodeling studies^[39]^. Therefore, further research should focus on adapting existing 3D registration methods to accommodate the unique deformation patterns of microvessels while preserving undetectable surface changes during this process.

Regarding the time-series surface mesh data, we propose our MAGIN method for vertex correspondence searching and deformation vector calculation. We assume the main axis of parent vessel internal surface keep static, similar to the static parent vessels in simulation models^[40,41]^, and measure the external surface deformation. A key advantage of the Gaussian radial basis function method is its flexibility in adjusting control point intervals and density. This enabled us to use parent vessel-sprout features^[14]^, construct a well-refined mesh specifically on the sprout spots, and achieve delicate yet large-scale local deformation without increasing overall computational consumption. The hierarchical registration strategy, guided by morphological information and developed in this study, achieved good registration results when dealing with large-scale irregular deformation patterns, such as the sprout region α highlighted in Figure 2c of Section 2.2. We also believe this method can be adapted to other surface segmentation methods, such as shape diameter-based segmentation, to deal with more complex situations. Because all deformations were measured as vectors, this registration-based deformation measurement allowed us to investigate the local deformation quantity, direction, and heterogeneity of angiogenic morphological changes in different areas.

As a hallmark of angiogenesis, angiogenic sprout areas had a larger mean deformation distance than parent vessel areas, and the majority of sprout vertices has local growth, as intuitively expected (Figure 3b and Figure S4). Meanwhile, parent vessel vertices had smaller deformation distances, and a relatively larger proportion of these vertices had temporal local regression (Figure 3b and Figure S4). This is consistent with the findings of in vivo angiogenesis studies on static parent vessels and active angiogenic sprouts^[7,42]^, further demonstrating that angiogenic morphogenesis is led by sprouts.

When we further analyzed the spatial distribution of deformation states, we found that many sprouts had local regression at their root side (Figure 3d), where we previously observed this type of pericyte attachment^[14]^. Because diameter enlargement is considered to negatively impact sprout outgrowth^[42,43]^, several factors, including the perturbation of vascular endothelial growth factor receptor (VEGFR)^[44]^, excessive vascular endothelial growth factor (VEGF) environment^[45]^, and shear stress signal^[42]^, have been shown to cause diameter enlargement, we hypothesized that root regression is related to the vascular stabilization function of pericytes. However, even though we can directly observe local sprout diameter control in some time-lapse videos^[46]^, to our knowledge, no studies discuss the sprout root area regression during morphogenesis and its possible mechanism. The root area is the junction between a sprout and its parent vessel; understanding the dynamics of sprout root diameter regulation can contribute to effective in vitro microvessel network construction and regenerative treatment strategies.

Both our previous study and the current one used a pericyte-embedded collagen gel to replicate the stem cell implantation microenvironment and subsequent therapeutic angiogenesis process. In contrast to typical in vivo angiogenesis, where pericytes are initially attached to the backside mature vessel, pericytes in this model were recruited to the vessel. Our previous study^[14]^ not only quantitatively measured the curvature-oriented behavior and angiogenesis regulation functions of pericytes but also mentioned three issues that were difficult to resolve without using the temporal method: 1) Does pericyte migration occur in a directed or random manner? 2) Do pericytes remain at the concave/convex regions, or do they migrate away over time? 3) Can pericytes still promote sprout morphological maturation (e.g., elongation, surface smoothing) if they are not present in the site? Answering these questions will help us gain a better understanding of pericyte dynamics and their potential in regenerative treatment.

In session 2.5, we answered our initial questions and explored their connection to microvessel morphogenesis. We first reviewed the pericyte coverage change to answer the first two questions. The slightly increased coverage rate across the entire microvessel and the significantly increased proportion of sprout area within the total covered area together demonstrated the bias during pericyte migration and settlement (Figure 5a, 5b & 5c). This suggests that pericytes sensed their local environment and selectively chose the sprouts, rather than randomly floating within the gel. The pericyte coverage state preservation ratio of less than 50%, presented in Figure 5c, showed that pericytes tended to change their position temporally. However, the difference between sprout vertex and the general level showed that settled pericytes were more likely to choose the sprout area than elsewhere, indirectly demonstrating the tacticity and the existence of pericyte-EC crosstalk.

We systematically evaluated pericytes’ influence at the single vertex level (Figure 6a & 6b), the sprout region level (Figure 6c & 6d), and the whole sprout level (Figure 7a & 7b). We found that pericyte-covered vertices had unique local deformation characteristics, including a small deformation distance and a high growth rate (Figure 6a & 6b). The small deformation distance may correlate with EC junction strengthening and vascular barrier stabilization in the presence of pericytes^[24]^. The high growth rate indicated that pericytes can initiate sprout formation. Furthermore, when we incorporated local geometrical information into the analysis, we observed a difference in the distribution of growth and regression states between pericyte-covered and -uncovered sprouts (Figure 6c & 6d). The heterogeneous angiogenic dynamics of pericyte-covered sprouts suggest the existence of a signaling pathway that may be triggered only by direct pericyte-EC contact, ultimately contributing to increased heterogeneity.

N1ICD is an activation marker of the Notch signaling pathway^[47,48]^, which is essential for directing angiogenesis and vascularization, and is particularly involved in pericyte-EC interplay^[49]^ and vessel diameter control^[50]^. Previous studies have shown that pericytes activate the EC Notch signaling pathway, stabilize the endothelium, and enhance the vascular barrier^[49]^. Here, we found local Notch activation in the pericyte-covered root area, whereas in sprouts without pericyte coverage, the Notch-activated areas were more evenly distributed. High MMP-1 expression has been shown to be related to collagen degradation, matrix scaffold breakdown^[51]^, and subsequent vascular structure regression^[52]^. Another study reported that MMP-1 also induces vasoconstriction via protease-activated receptor-1 (PAR-1)^[53]^. However, MMP-1 can also enhance microvessel angiogenesis^[54]^ and has been found to up-regulate vascular endothelial growth factor receptor-2 (VEGFR2)^[55]^, which typically has higher expression levels in the front tip cells of the sprout^[56]^. In this study, we used MMP-1 expression levels to assess sprout regression and growth. Our results showed a heterogeneous distribution pattern of MMP-1 signal, with the highest expression at the root, almost no signal in the central area, and a relatively low level at the tip (Figure 6e & 6f). The low expression in the central area is consistent with the MMP-1 inhibition effect of pericyte-EC coculture reported previously^[57]^. An overall down-regulation of MMP-1 was also observed in our direct fluorescence intensity comparison between monoculture and coculture sprouts (Figure S7). Intriguingly, the two high expression level zones, the tip and root, coincide with the preferred pericyte settlement positions^[14]^, indicating that the pericyte-EC crosstalk pattern is adjusted according to the local morphology. To our knowledge, this is the first paper to discuss MMP1 distribution in relation to pericyte coverage and local angiogenic morphogenesis.

We finally discussed the overall morphology change of pericyte-covered and -uncovered sprouts (Figure 7a & 7b). We found that both the pericyte-covered and -uncovered sprouts became smoother and more elongated; however, pericyte-covered sprouts consistently had greater surface smoothness and length than uncovered sprouts, which once again suggests that pericyte-EC contact enhances the sprout maturation process and that this effect may not be replaced just by the long-distance cell–cell crosstalk.

In the future, the feasibility of the MAGIN method for angiogenesis measurement should be further verified. Adaptations for complex vascular structure analysis, such as dense vascular networks in organoids^[58]^, and for depth-dependent morphogenesis analysis, such as retinal angiogenesis^[16]^, are expected to be developed. Complex crosstalk mechanisms, such as secretion factors for distant interactions (paracrine) or membrane proteins for direct contact interactions (juxtacrine), will be considered.

### Experimental section

Supporting methods are available in the supporting information.

### Microscopy observations

Before observation, HUVECs were stained for 15 min with 20 μL of EGM-2 medium diluted with 20 μg/mL rhodamine-conjugated UEA-1 and then washed with fresh medium. Panoramic images of the microvessels were taken using an Axio Observer Z1 microscope (Carl Zeiss, Oberkochen, Germany) with a 20× objective lens (LD Plan-NeoFluar 20×/0.4 Korr Ph2 M27). A 12 × 2 image scanning tile with a 10% stitching overlap was used. Images of each chip were taken on the day after HUVEC seeding (Day 1) and five days after seeding (Day 5).

3D microvessel morphology was observed using a confocal laser scanning microscope (CLSM; LSM 700, Carl Zeiss, Oberkochen, Germany) with 20x objective lenses (Plan-Apochromat 20×/0.8 and LD Plan-NeoFluar 20×/0.4 Korr M27). Images were acquired in the Z-direction at 2 μm intervals and stored as image stacks. Each stack contained 126 pieces of images, ensuring that the parent vessel and most of the sprouts were visible. For each slice, eight pieces of 256 × 256 pixel images were taken at the same height using a tile scanning mode (horizontal × vertical: 4 × 2). Light at wavelengths of 488 and 555 nm was collected to visualize the vessel morphology (UEA-1, Rhodamine, red) and pericytes (GFP, green), respectively. Images were processed using ZEN 2 blue edition software (version 2.0.0.0, Carl Zeiss).

MMP-1, N1ICD, and CD31 were observed using a 40 × objective lens (LD C-Apochromat 40 ×/ 1.1 W Korr M27). 1024 × 1024 pixel images were acquired in the Z-direction at 2 μm intervals and stored as image stacks. The image stack included the sprout region of interest (ROI). Light at wavelengths of 408 nm, 488 nm, and 555 nm was collected to visualize the localization of N1ICD, MMP-1 (Alexa Fluor® 405, cyan), pericytes (GFP, green), and vessel morphology (CD31, Alexa Fluor® 555, red), respectively. Images were processed using ZEN 2 blue edition software and FIJI to create Z-stack maximum intensity projection images (MIPs).

### 3D mesh reconstruction

Both the red and green channels of 3D confocal microscopy images were treated with a 3 × 3 × 3 median filter to remove noise signals. Next, the de-noized images were processed using IMARIS (version 9.0.0, BitPlane, Zurich, Switzerland), a 3D image visualization and processing software. Images were binarized according to a built-in, self-adapting threshold algorithm, and the triangular surface meshes of microvessels and pericytes were created separately. Mesh components with fewer than 1000 vertices were removed before the final export.

### Mesh pre-alignment

Meshes from two consecutive days were paired for processing. The mesh from the earlier day was designated as the source data, and the mesh from the later day as the target data. The two meshes were manually overlapped, and some extra side parts were removed. We assumed the central line of the microvessel remained static and used the centroid and the central line of the internal surface mesh to align the source data to the target data. The vector from the source mesh centroid to the target mesh centroid was used as the translation vector, T. A rotation matrix, R, was calculated using Rodrigues’ rotation formula^[59]^ to align the normalized central line vector of the source vessel mesh with that of the target mesh (Figure S2). The rigid transformation, defined by translation vector T and rotation matrix R, was then applied to the external surface mesh of the source data. The vertex positions of the aligned source data served as the initial positions for subsequent vertex deformation calculations.

### Mesh registration

The deformation calculation was based on a Gaussian radial basis function method mentioned in a previous study^[26]^.

The registration process consists of coarse grid-controlled global deformation, fine grid-controlled sprout area deformation, and local mesh fitting. At the global deformation step, coordinates of the source and target vessel mesh vertices were extracted and placed into a 3D space. The bidirectional correspondences between the target and source vertices were used to eliminate the size difference between the two datasets and to provide extra displacement in the sprout position to wrap the source mesh to the target mesh. Thus, the augmented source vertex dataset 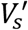 was made up of the original source vertices, denoted as 𝑉*_So_*, and the repeated vertices, 𝑉_𝑠𝑟_, which consist of the closest source vertices of each target vertex. Similarly, the augmented target vertex dataset 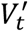 also contained the original vertices, 𝑉_𝑡𝑜_, and the repeated vertices, 𝑉_𝑡𝑟_. The deformation vector 𝑣_1_ from the augmented control point 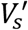 to augmented target control point 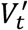 is given by E-1. This vector was used as the morphology constraint for the global deformation.

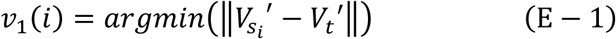

Then, a grid was constructed in the space, and the grid nodes near the source vertices were selected as the deformation control points. Grid nodes near the source vertices were selected as the deformation control points, 𝑉_𝑐1_. This correspondence between the augmented source vertex dataset 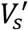 and deformation control points 𝑉*_c_* was defined through a Gaussian radial basis function, as shown in E-2 and E-3. A vertex can be affected by all the control points on the grid, while the weight of the transformation passage was based on the Euclidean distance 𝑟 , indicating that the vertices near the grid nodes had a larger deformation. ε was the width of the Gaussian function, which was set to be proportional to the average of the distance 𝑟 between all source vertices 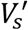 and all control points 𝑉_𝑐1_ ^[27,60]^.

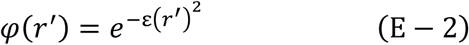

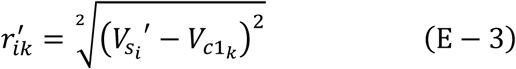

Combining the distance relations between augmented source points to all the control points mentioned in E-2 and E-3 and the relative position information between 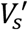 and 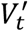, which was denoted as vector 𝑣_1_ in formula E-1. This problem can be transformed into a least squares problem to find the deformation matrix 𝑑 of control points, to minimize 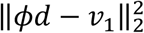, where 𝜙 is the deformation transfer matrix. An optimization step with a Tikhonov term was performed to conduct this translation from the vector 𝑣 and the displacement 𝑑, while minimizing the difference and preventing irregular mesh stretching, as shown in E-4^[61]^. The parameter 𝜆 was empirically defined as 0.001 in this step.

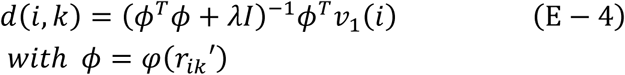

Subsequently, the deformation 𝑑 at each control point was transferred back to the original source vertices set 𝑉_𝑠_, again through the Gaussian radial basis function, as shown in E-5.

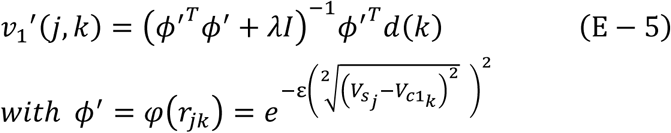

This global deformation process was iterated from coarse to fine, along with a stepwise decrease of the deformation grid voxel cube side length and the width ε of the Gaussian radial basis function.

Sprout areas on the target mesh were always larger and contained more vertices than their corresponding area on the source mesh. In such cases, an augmentation of the source sprout vertex set was conducted to mimic the possible nearby area pulling phenomenon during sprout growth and elongation. A set of control points 𝑉_𝑐1_^𝑠𝑝𝑟𝑜𝑢𝑡^ close to source mesh sprout vertices 𝑉_𝑠_ ^𝑠𝑝𝑟𝑜𝑢𝑡^ was selected from 𝑉_𝑐1_ (E-6). Then, the source vertices whose closest control point was among 𝑉_𝑐1_^𝑠𝑝𝑟𝑜𝑢𝑡^were extracted from 𝑉_𝑠_ as the augmented sprout vertices set 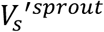 (E-7). Finally, the control points of the fine grid 𝑉_𝑐2_ were extracted by searching the nearby grid nodes for the augmented sprout vertices set 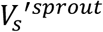 (E-8). The parameter 𝜆 was empirically defined as 0.5 in this step.

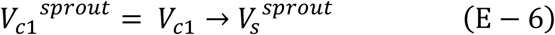

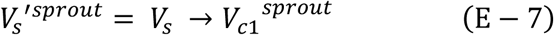

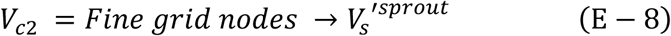

The calculation process was similar to the coarse-grid-controlled deformation, but only the difference in the sprout area (𝑣_2_), as shown in E-9, was considered. A distance filter, ‖𝑣_2_(𝑖)‖ should be less than 3 times the current control point interval, and a normal direction filter, 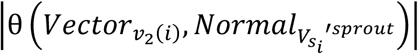 should be less than 60 degrees, was applied to prevent mispairing. After calculating the deformation matrix 𝑑 of control points, the deformation vector was passed back to the whole source mesh to ensure a smooth deformation.

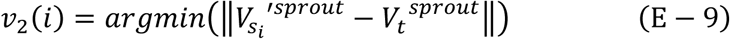

Once the side length of a triangle face becomes 3 times larger than the mean side length of the whole mesh and the aspect ratio was larger than 10, this triangle face is split into four triangles through the midpoint-split method to prevent incorrect deformation. All the new vertices produced by face-splitting events and the vertices of their parent triangle face were stored.

After the grid-controlled deformation, a local mesh fitting was applied to every vertex. For each vertex in the source data, about 15 nearby vertices in the source data and 3 nearby vertices in the target data were selected. A Procrustes analysis with a scaling and reflection restriction was performed to calculate the deformation matrix from nearby source vertices to nearby target vertices. Then, the deformation is passed back to the selected source vertex according to a weight coefficient 𝑊. This local deformation was also repeated 10 times.

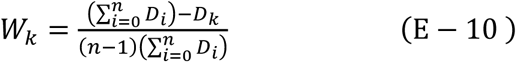

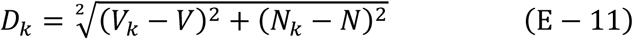

where 𝑉 and 𝑉_𝑘_ are the coordinates of the selected vertex and the k-th nearby vertex of the selected vertex, respectively. 𝑁 and 𝑁_𝑘_ are the vertex normal vectors of the selected vertex and the k-th nearby vertex of the selected vertex, respectively^[26,28]^.

The traditional iterative closest point (ICP) registration algorithm and a B-spline grid-based non-rigid transformation have also been used to align the 3D meshes from two consecutive days for comparison. Manually registered raw vessel mesh data was processed by ICP (rigid), B-spline, and radial basis Gaussian function-based registration (non-rigid) methods described in this section. Results show that the Gaussian radial basis function-based registration method achieved the best result compared with the other two methods, as shown in the supplementary file (Figure S4).

### Deformation vector and deformation distance measurement

For shape mapping, we assumed the source vessel mesh closely approximated the target microvessel mesh after deformation. For each target microvessel mesh vertex, its nearest vertex on the source vessel mesh was identified and stored as its corresponding point. The deformation vector was defined as the vector originating from the corresponding point’s original position on the source mesh before non-rigid registration and ending at the target vertex. The Euclidean length of this vector was used as the deformation distance. If a corresponding point originated from face splitting during deformation, its original position was considered the centroid of its two parent vertices. This search was repeated if the parent vertices were also generated from face splitting.

### Statistics

Microvessel meshes on days 3-5 were reconstructed from seven independent experiments, and the deformation of individual microvessels at different time periods was treated as independent datasets to perform statistical comparisons. Two-sample Kolmogorov–Smirnov tests were used to evaluate the differences in the distribution of angles between deformation and normal vectors for parent vessel and sprout vertices (Figure 3d), the spatial distribution of growth/regression states (Figure 6c & 6d), and the differences in the distribution of angles between deformation and normal vectors for pericyte-covered and -uncovered vertices (Figure 6b). Seven different microvessels were considered as time-series datasets in the temporal pericyte coverage rate analysis. Differences between the time points were evaluated using a paired permutation test (Figure 5). Welch’s t-test was used to evaluate the difference in mean deformation distance between pericyte-uncovered and pericyte-covered vertices (Figure 6a). Comparisons of morphological parameters between pericyte-uncovered and pericyte-covered sprouts, and comparisons between the previous day and the subsequent day, were evaluated using paired and unpaired permutation test, respectively (Figure 7a & 7b). All statistical differences were considered significant when p-values were < 0.05.

## Acknowledgements

H.Z. acknowledges the financial support of Otsuka Toshimi Scholarship Foundation, and the fellowship from JST SPRING (Grant Number JPMJSP2108). The authors also thank Prof. Marie Shinohara (The University of Tokyo) and Jules Edwards (LAAS-CNRS) for their constructive comments.

## Author Contributions

H.Z., T.S., and Y.T.M. designed the study. H.Z. performed research. H.Z. and T.S. analyzed data. J.K. prepared cells. H.Z. and T.S. wrote the manuscript. Y.T.M. supervised the research. All authors have reviewed the manuscript and approved its contents.

## Data Availability Statement

The data that support the finding of this study are available from the corresponding author upon reasonable request.

