## Supporting_info_SI_method_figures for "Spatiotemporal analysis of pericyte-induced dynamics in nascent angiogenic morphogenesis and heterogeneity using a microvessel-on-a-chip platform"

Yukiko T. Matsunaga, Ph. D.

Institute of Industrial Science, The University of Tokyo 4-6-1 Komaba, Meguro-ku, Tokyo 153- 8505, Japan. Tel.: +81-3-5452–6470; Fax: +81-3-5452-6471;

**Table of Contents**


**1. Supplementary Method**

**2. Supplementary Figures**

Figure S1. Step results of external mesh extraction.

Figure S2. PCA axis-based mesh pre-alignment.

Figure S3. Schematic illustration of different registration methods and their effects on distance-based shape mapping and correspondence searching.

Figure S4. Comparison among iterative closest point (ICP), B-spline grid and Gaussian radial basis function registration methods.

Figure S5. Measurement and comparison of deformation distance between parent vessels and sprouts.

Figure S6. Probability distribution histograms of relative positions for: (a) pericyte-covered and (b) pericyte-uncovered sprout vertices, and (c) pericyte-covered and (d) pericyte-uncovered parent vessel vertices.

Figure S7. Fluorescence intensity comparison between monoculture and coculture chips.

**3. Supplementary Movie**

Movie S1. Linear interpolation animation of an angiogenic sprout deformation.

### 1. Supplementary method

#### Reagents and materials

Type I-A collagen (acid-extracted collagen isolated from porcine tendons, 3 mg/mL, pH 3) was purchased from Nitta Gelatin Inc. (Osaka, Japan). Acupuncture needles (J type, No.02, 200 μm diameter and 30-mm length) were purchased from Seirin Co. Ltd. (Shizuoka, Japan). 10x phosphate-buffered saline (-) (PBS) was purchased from FUJIFILM Wako Pure Chemical Industries Ltd. (Osaka, Japan). Dextran from *Leuconostoc* spp. (Mr 450,000-650,000) and Hanks' Balanced Salt Solution were purchased from Sigma-Aldrich Co. LLC. (St. Louis, MO, USA). Rhodamine-conjugated *Ulex europaeus* Agglutinin 1 lectin (UEA-1) was purchased from Vector Laboratories (Burlingame, CA, USA). Water used in the experiment was collected from a Milli-Q system (Darmstadt, Germany) and then sterilized using an autoclave.

Polydimethylsiloxane (PDMS) chips for in-vitro microvessel fabrication were the same as those reported in our previous studies. Each chip has a cuboid chamber (2 mm × 6 mm × 2 mm: width × length × height), two reversed truncated cone-shaped wells (top circle diameter: 3 mm, bottom circle diameter: 2 mm, height: 6 mm), and a cut-through hole connecting them at the bottom.

#### Cell source and cell culture

Human umbilical vein endothelial cells (HUVECs) were purchased from Lonza (Basel, Switzerland). Pericytes expressing the ephrin type-A receptor 7 (EphA7) gene and green fluorescent protein (GFP) were isolated from the capillaries of subcutaneous adipose tissue in mice.

Endothelial cell growth medium-2 BulletKit (EGM-2) was purchased from Lonza (Basel, Switzerland). Vascular endothelial growth factor-A165 (VEGF-A165) was purchased from FUJIFILM Wako (Osaka, Japan). EGM-2 was supplemented with 15 ng/mL VEGF-A165 for immunofluorescence staining chips. Gibco™ Dulbecco's modified eagle medium (DMEM) (low glucose, with GlutaMAX™ supplement and pyruvate) was purchased from ThermoFisher Scientific (Waltham, MA, USA). The pericyte culture medium was prepared by mixing DMEM with the following components: 4% fetal bovine serum (purchased from Biosera, Nuaille, France), insulin-transferrin-selenium-sodium pyruvate (ITS-A) (100×) (purchased from Gibco, Thermo Fisher Scientific, Waltham, MA, USA), penicillin-streptomycin (purchased from FUJIFILM Wako, Osaka, Japan), 0.05% bovine serum albumin (BSA) (purchased from Sigma-Aldrich, St. Louis, MO, USA), 20 ng/mL recombinant murine epidermal growth factor, and 20 ng/mL recombinant murine fibroblast growth factor (FGF)-basic (PeproTech, New Jersey, USA).

For the experiments, frozen HUVECs and pericytes were thawed in EGM-2 and modified DMEM medium, respectively. HUVECs (passages 4-7) and pericytes (passages 10-12) were used in this study. Thawed cells were then seeded in a 6 cm diameter dish and cultured in a 37°C, 5% $CO_2$ incubator for amplification. After two days of culture, the HUVECs and pericytes were washed with 1x PBS solution and treated with 0.25% trypsin-ethylenediaminetetraacetic acid (EDTA) solution for 3 minutes in a 37°C, 5% $CO_2$ incubator. The cells were then harvested using EGM-2 and modified DMEM medium respectively, for use in subsequent experiments.

**Microvessel (MV) chip fabrication**

PDMS chips and acupuncture needles were first cleaned in an $O_2$ plasma cleaner for 5 min and then treated with 3-aminopropyl trimethoxysilane (Sigma-Aldrich, St. Louis, MO, USA) vapor in a vacuum chamber for 30 min. The chips were then baked in an oven at 70°C for 10 min to remove the excess 3-aminopropyl trimethoxysilane. Next, 50 μL of 2.5% glutaraldehyde was added to the chip's central chamber for 1 min and then removed. After washing 10 times with sterile water, the chips were sterilized under UV light and placed on a clean bench to dry.

The cell concentration of the pericyte suspension was first counted using a hemacytometer. Before applying the collagen gel, the necessary suspension volume containing $2 \times 10^4$ pericytes was calculated and collected in a separate tube. Collagen gel was prepared by mixing collagen I-A, Hank's Balanced Salt Solution, and collagen buffer in an 8:1:1 ratio. The prepared collagen was then thoroughly mixed with the centrifuged pericyte precipitate and applied to the central cubic chamber of the PDMS chip. A 200-μm diameter BSA/PBS-coated acupuncture needle was inserted from one side before gel polymerization.

After needle insertion, the microvessel chips were returned to the incubator for 45 min to allow the gel to cure. Then, 1 mL of warmed EGM-2 medium was added to each chip, and the whole chip was returned to the incubator for 24 hours. The next day, the acupuncture needle was removed, leaving a channel within the gel. Both sides of the chip's bottom hole were sealed with two short stainless-steel needles. Beforehand, 10 μL of a 1 mg/mL fibronectin solution was added to both wells to coat the central lumen. Next, 4 μL of HUVEC suspension at a concentration of $1 \times 10^7$ cells/mL was added to the side wells. Excess HUVECs were washed out, and fresh EGM-2 was added to the chip. The culture medium was changed daily.

**Immunofluorescence**

After a 5-day culture, the chips were fixed with a 4% PFA solution and washed with PBS. Then, the chips were permeabilized with 0.5% Triton X-100 in PBS solution, washed with 0.1% PBST, quenched with 750 mM Tris, and blocked with 1% BSA in PBS solution for 2 hours at 25°C. After blocking, a mixture of 1:50 diluted rabbit monoclonal anti-human MMP-1 antibody (ab52631, Abcam, Cambridge, UK) and 1:200 diluted mouse anti-human CD31 antibody (M0823, DAKO, Santa Clara, CA, US), or a mixture of 1:100 diluted sheep anti-human Notch-1 intracellular domain polyclonal antibody (AF3647, R&D systems, Minneapolis, MN, US) and 1:200 diluted mouse anti-human CD31 antibody was added to the chips. After overnight storage in a 4°C fridge and subsequent primary antibody washout, a secondary mixture of either 1:300 diluted donkey anti-rabbit IgG H&L (Alexa Fluor® 405) (ab175651, Abcam, Cambridge, UK) and 1:250 diluted sheep anti-mouse IgG H&L (Alexa Fluor® 555) (A21422, Invitrogen, Thermo Fisher Scientific, Waltham, MA, USA) or 1:300 diluted donkey anti-sheep IgG H&L (Alexa Fluor® 405) (ab175676, Abcam, Cambridge, UK) and 1:250 diluted sheep anti-mouse IgG H&L (Alexa Fluor® 555) was added to the chips, respectively. The chips were wrapped in aluminum foil, placed at room temperature, and then washed with PBST.

**Mesh division**

Microvessel meshes reconstructed from the CLSM images were imported into MATLAB (Natick, MA, USA). The microvessel meshes were then segmented into external and internal surfaces. We first defined the first principal component analysis (PCA) loading of the surface mesh vertices as the microvessel central line. The centroid position and the normal vector of each mesh face were calculated. A perpendicular line was then drawn from the face centroid to the central line, and the angle between the face normal and this perpendicular line was computed. Faces with angles less than 75° were classified as internal meshes, and those with angles greater than 105° were classified as external mesges. Faces with angles between 75° and 105° were classified based on mesh connectivity. A recursion function was used to iteratively search the neighbors of those faces, until encountering an "internal" or "external" face. The same label was then temporarily given to the pending faces to complete the coarse classification. A refinement step was then performed to minimize potential misclassifications. The largest connected components in the "internal" and "external" mesh groups were identified. The distribution of distances between the surface faces and the microvessel central line was calculated for both meshes, along with their respective normal distribution fits. The 95% confidence intervals of these normal distributions were used to reclassify other small-sized connected components within the “internal” and “external” mesh groups. The internal vessel surface meshes were used to calculate the vessel’s central line for the subsequent registration steps. The external vessel surface meshes were used for parent vessel-sprout classification, non-rigid mesh registration, and deformation measurement. A visualized mesh cropping process can be found in supplementary file Figure S1.

**Sprout extraction**

The sprout extraction step was modified from the method we proposed in our previous study (T. Sano et al., ***Stem Cell Research & Therapy*** 2022, 13, 532.). For each mesh, a rotation matrix was calculated to align the central line of the internal surface mesh with the positive x-axis (1, 0, 0). This matrix was then applied to the corresponding external surface mesh originated from the same raw surface mesh, ensuring the longitudinal axis of the vessel mesh was perpendicular to the y-z plane. The external surface mesh was then converted into a voxel volume object at the same resolution as the raw z-stack image. This volume object was processed using a 3D fast Fourier transform. A low-pass filter was applied to retain only areas without frequent density changes along the x-axis, which typically corresponded to the parent vessel region of the microvessel. The remaining parts with areas larger than 100 voxels after parent vessel extraction were classified as sprouts. The extracted parent vessel and sprout volume objects served as masks for classifying the mesh vertices. Vertices within the sprout component were identified as sprout vertices and assigned a sprout number according to sprout mask mesh connectivity.

**Pericyte coverage analysis**

The vessel mesh and the pericyte mesh, generated from the same confocal image data, were used to calculate pericyte coverage. First, pericyte meshes from two consecutive days were imported into MATLAB software, and the same transformation used in “Mesh coarse alignment” was applied to the pericyte mesh from the previous day. Then, pericyte meshes were placed into a 1024 × 512 × 126 logical matrix and transformed into a discrete volume object using a MATLAB community function developed by Dirk-Jan Kroon (https://www.mathworks.com/matlabcentral/fileexchange/24086-polygon2voxel). After applying MATLAB’s built-in image dilation, erosion, and hollow filling functions, the inner hollow of the volume object were filled. The vessel mesh vertices were then placed into the matrix. A vertex placed in a voxel where the signal was true was judged to be a pericyte-covered vertex. Otherwise, it was judged to be a pericyte-uncovered vertex.

**Sprout correspondence searching and length measurement**

For each sprout component obtained from the “sprout extraction” step, the centroid of sprout mesh vertices, the first three PCA loading vectors, and the bounding box along the first three PCA loading vector directions were measured. To search for the correspondence between two sprouts at two different time points, a virtual vector was defined, starting from the centroid of the sprout on the previous day and pointing to the Euclidean coordinates of the sprout centroid on the subsequent day. The projections of this virtual vector onto the directions of the first three PCA loading vectors of the previous day’s sprout were then calculated. If the projection in one direction was included in the bounding box, it was counted as one point. The same process was performed in reverse, from the subsequent day’s sprout centroid to the previous day’s sprout centroid, allowing one sprout pair to obtain a maximum of six points in total. The number of points was considered a geometrical correlation index, and selected the sprout pair with corresponding point $\geq 3$.

A sprout could correspond to more than one sprout on the other day. Sprout pairs that had one and only one corresponding sprout on the other day were specifically selected for temporal morphology change measurements in session 2.5. The STD of the Gaussian curvature of the sprout surface mesh was calculated using a function from an open-source MATLAB toolbox developed by Gabriel Peyre (https://github.com/gpeyre/matlab-toolboxes), and the sprout length was defined as the maximum difference after projecting all vertices onto the first PCA loading vectors.

### 2. Supplementary Figures

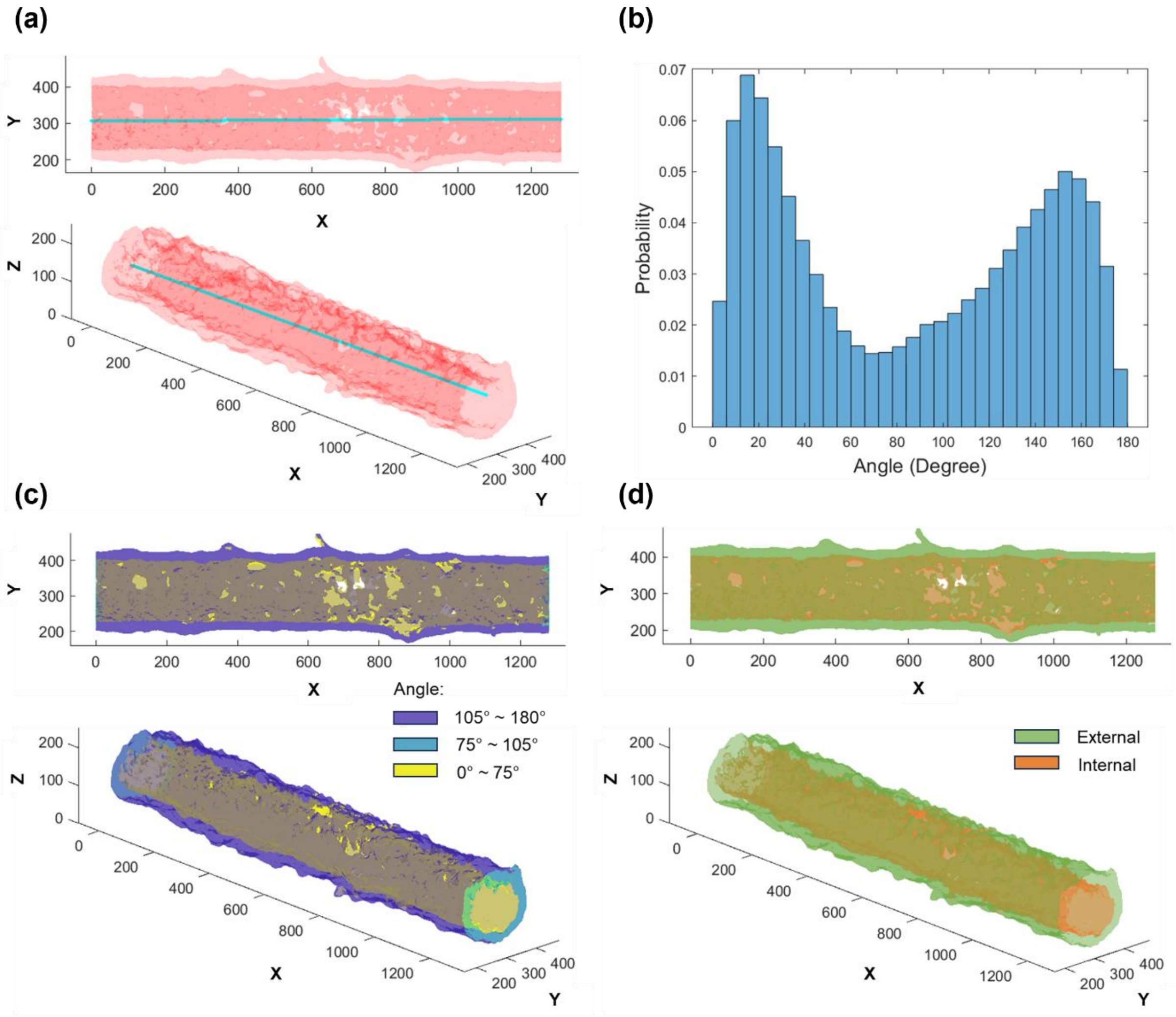


**Figure S1. Step results of external mesh extraction. (a)** The first loading axis obtained from a principal component analysis (PCA) of the microvessel surface vertex coordinates, was defined as the microvessel centerline. **(b)** A histogram shows the calculated angles between the face normal vectors and the centerline, defined from each face centroid to the first loading of the PCA. **(c)** Based on these angles, meshes were coarsely classified as extraluminal surfaces (purple), intraluminal surfaces (yellow), or image boundary surfaces (green). **(d)** Classification results were refined using mesh connectivity information.

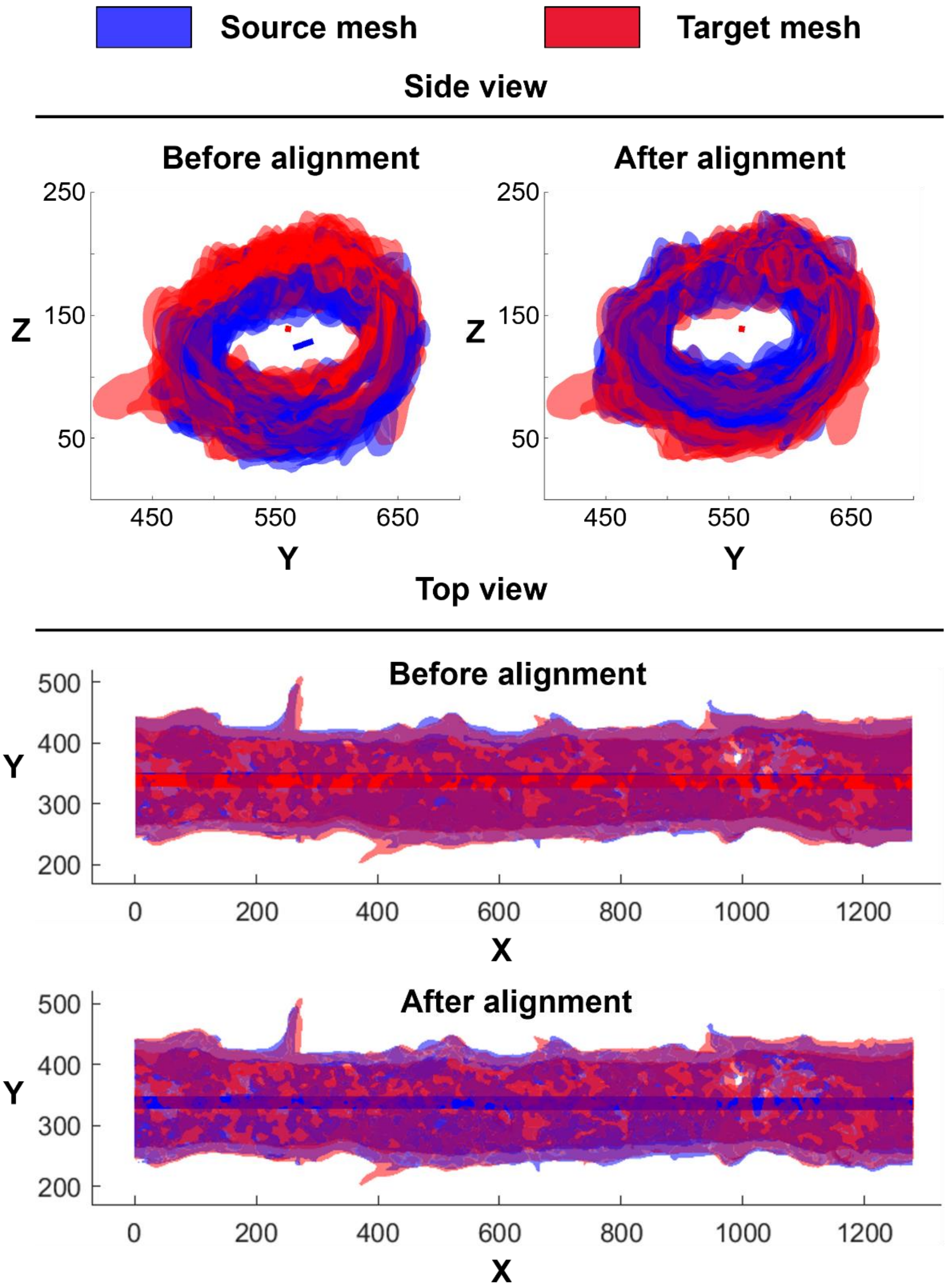


**Figure S2. PCA axis-based mesh pre-alignment.** The surface mesh reconstructed from day X data was rotated to let its PCA first axis of the internal surface be aligned with that of mesh reconstructed from day X+1 data. A representative surface mesh set from day 4 and day 5 were presented as “source mesh” and “target mesh.” Unit: μm.

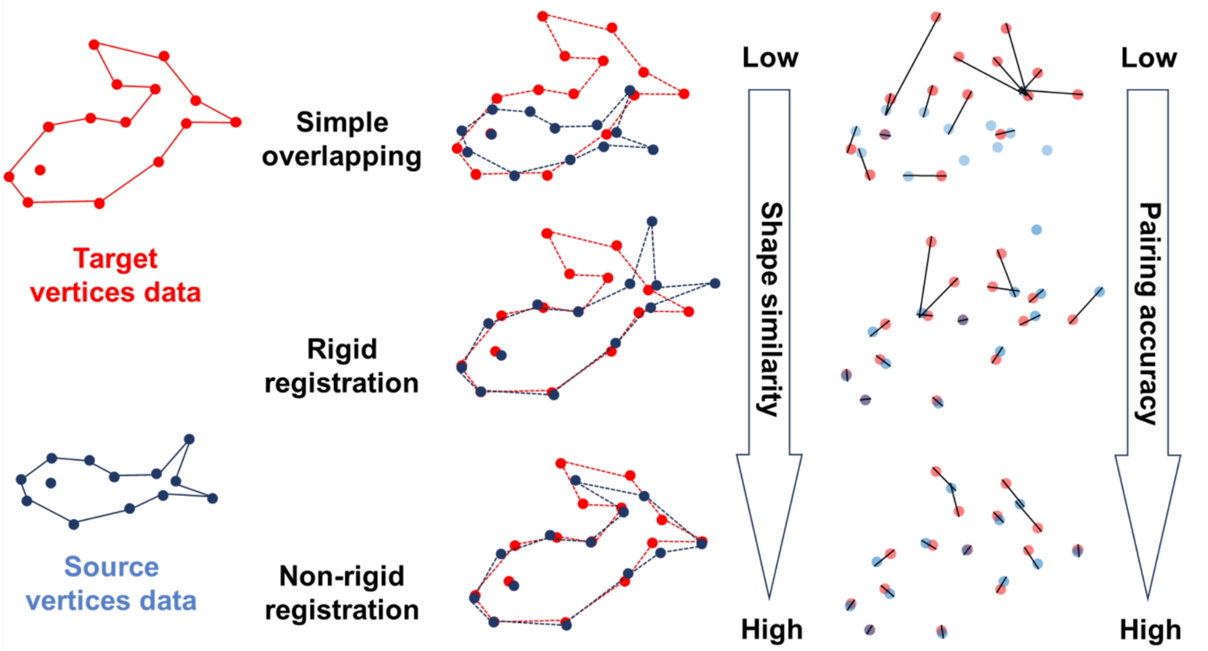


**Figure S3. Schematic illustration of different registration methods and their effects on distance-based shape mapping and correspondence searching.** Non-rigid registration allows for distinct transformations at different positions, facilitating more accurate vertex correspondence searching.

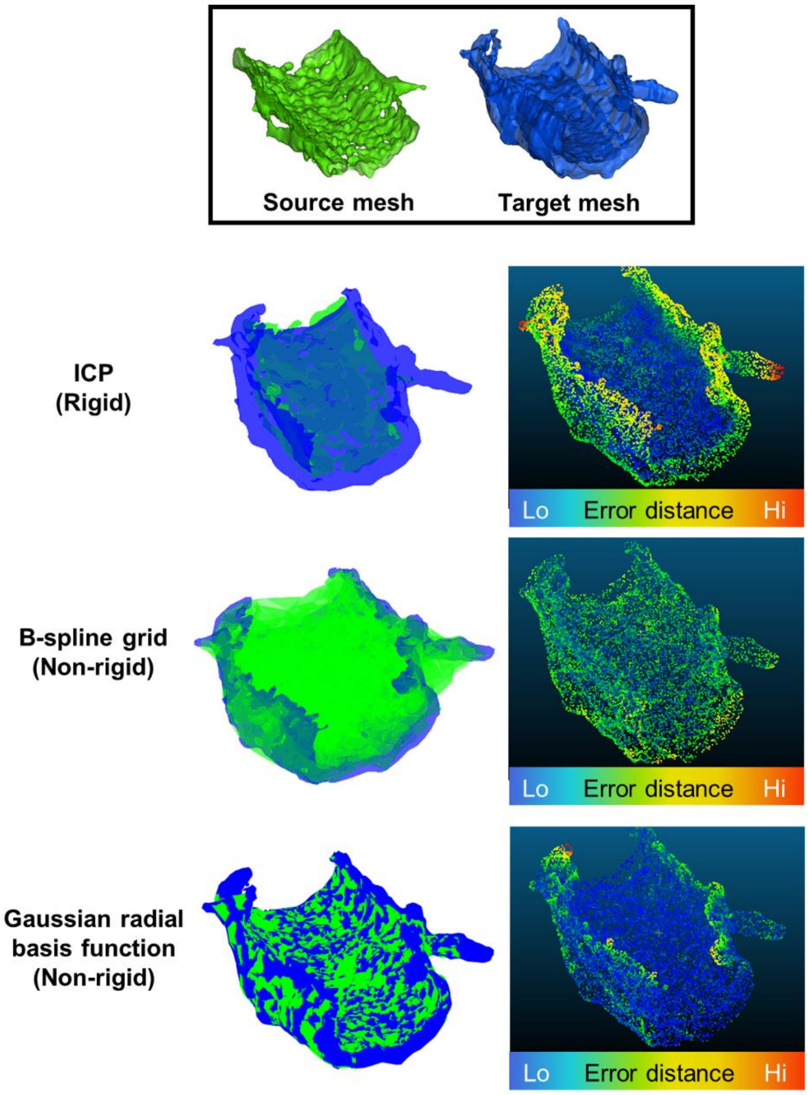


**Figure S4. Comparison among iterative closest point (ICP), B-spline grid and Gaussian radial basis function registration methods.** The source mesh, after undergoing Gaussian radial basis function-based registration, conforms to the target mesh more accurately than with other methods. The error distance was calculated as the Euclidean distance from each target mesh vertex to its nearest corresponding vertex on the source mesh. B-spline grid registration was performed using a MATLAB community function developed by Dirk-Jan Kroon (https://ww2.mathworks.cn/matlabcentral/fileexchange/20057-b-spline-grid-image-and-point-based-registration). Error distance was evaluated and visualized using CloudCompare software (https://www.danielgm.net/cc/).

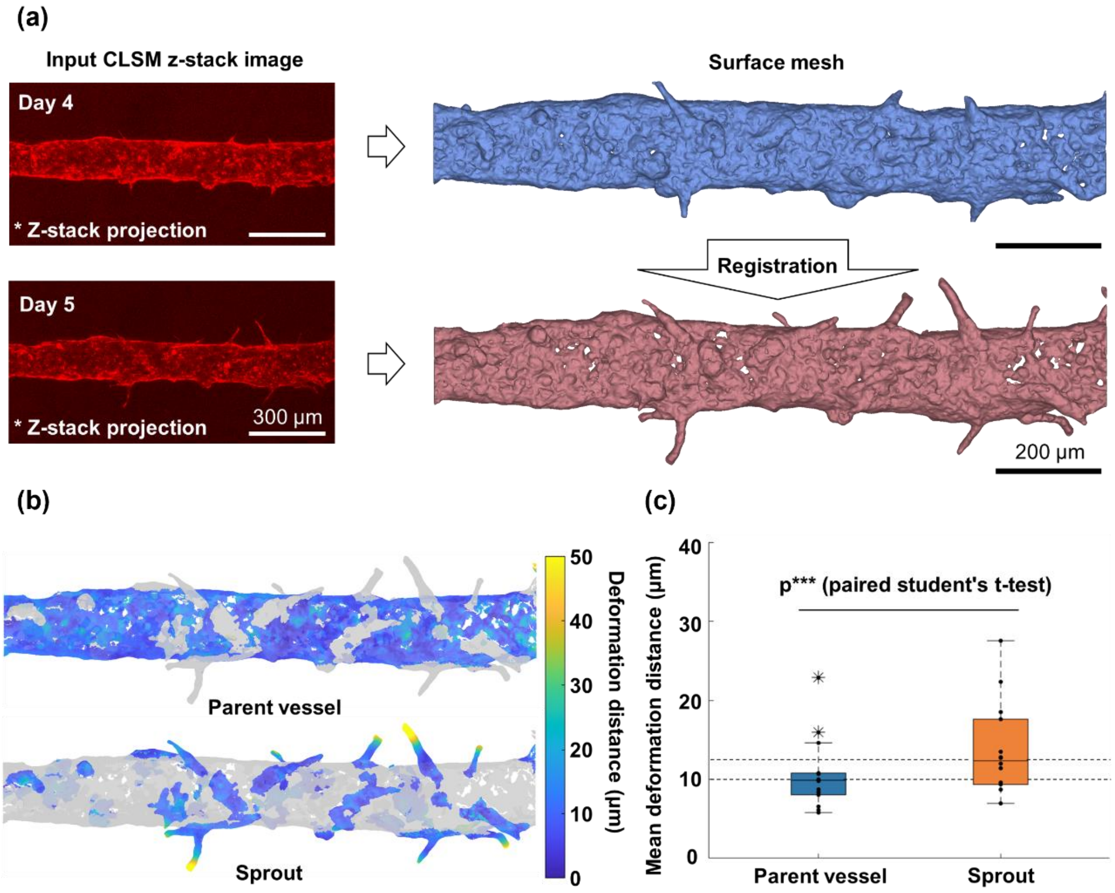


**Figure S5. Measurement and comparison of deformation distance between parent vessels and sprouts. (a)** Schematic illustration of representative 3D microvessel surface meshes reconstructed from time-series Z-stack confocal microscope images taken on day 4 and day 5. **(b)** Deformation colormap of the parent vessel and sprout regions on the day 5 mesh. Vertex deformation distance was calculated using vertex correspondence, determined after registration and segmentation of the entire day 5 structure into parent vessel and sprout regions. Deformation distance was defined as the Euclidean length of the vector from a source mesh vertex to its corresponding vertex on the target mesh. Scale bar: 300 μm. **(c)** Distribution of mean deformation distances for vertices in the parent vessel and sprout regions. Data from two time periods (day 3 to day 4 and day 4 to day 5) across seven samples were combined to generate the histogram. The median deformation distance for the sprout region (12.346 μm) was significantly greater than that for the parent vessel region (9.884 μm). Asterisks represent statistical outliers. p*** indicates $p \leq 0.001$.

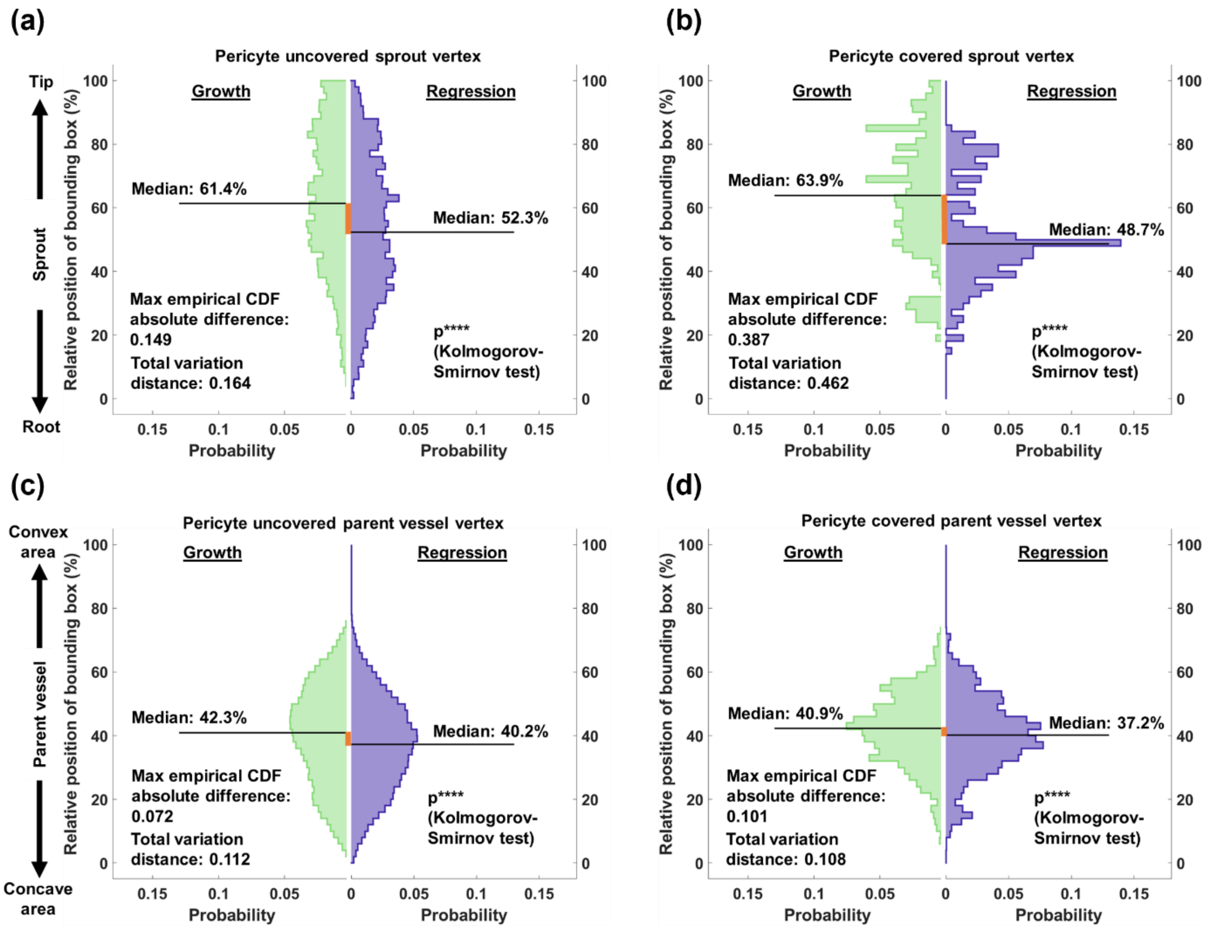


**Figure S6. Probability distribution histograms of relative positions for: (a) pericyte-covered and (b) pericyte-uncovered sprout vertices, and (c) pericyte-covered and (d) pericyte-uncovered parent vessel vertices.** Histograms were generated separately for growth and regression vertices.

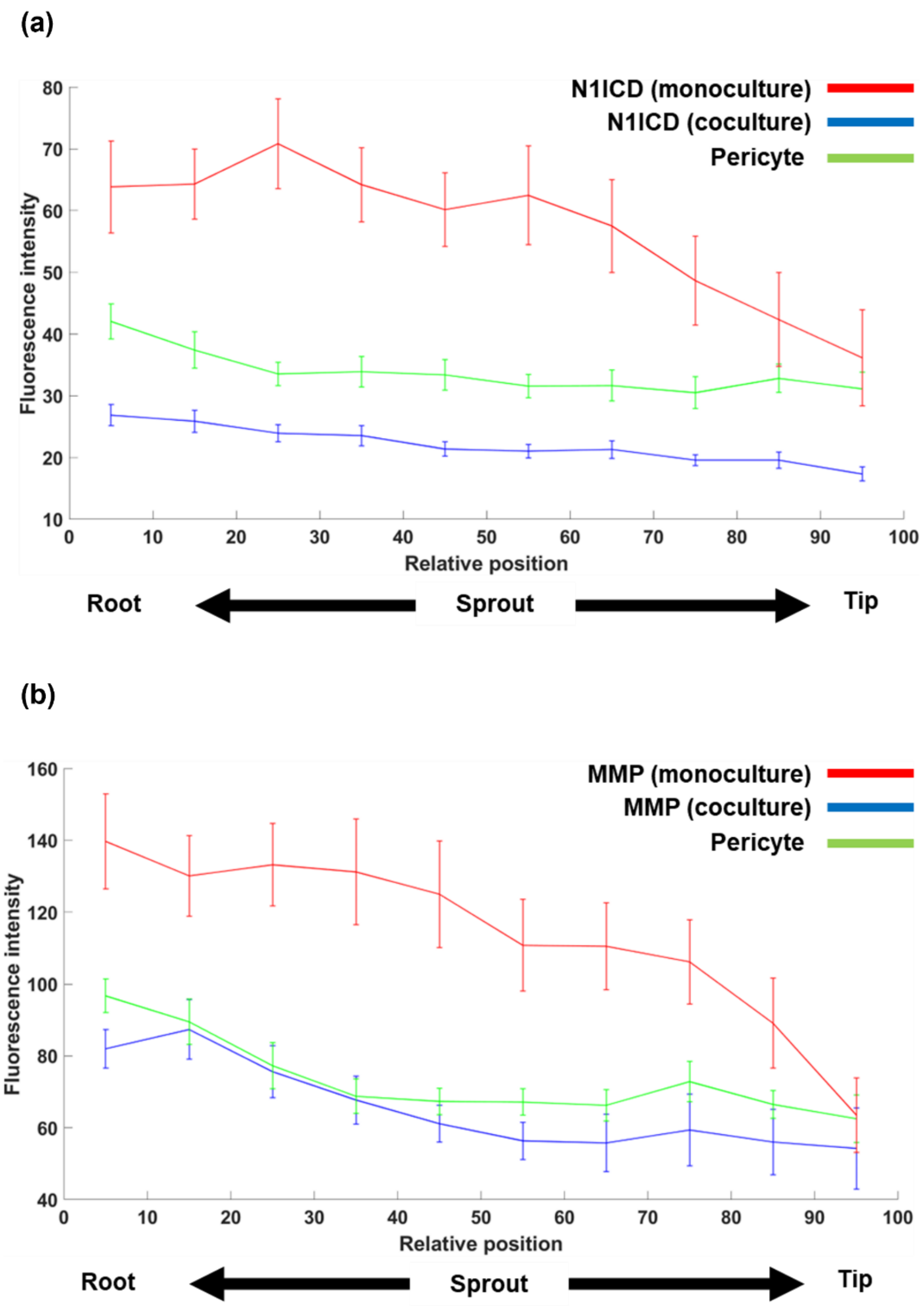


**Figure S7. Fluorescence intensity comparison between monoculture and coculture chips. (a)** Intensity comparison of Notch-1 intracellular domain (N1ICD) levels in mono-culture sprouts, N1ICD levels in co-culture sprouts, and green fluorescent protein (GFP) levels in pericytes. **(b)** Intensity comparison of matrix metalloproteinases-1 (MMP-1) levels in mono-culture sprouts, MMP-1 levels in co-culture sprouts, and GFP levels in pericytes. Error bars represent standard error.

### 3. Supplementary Movie

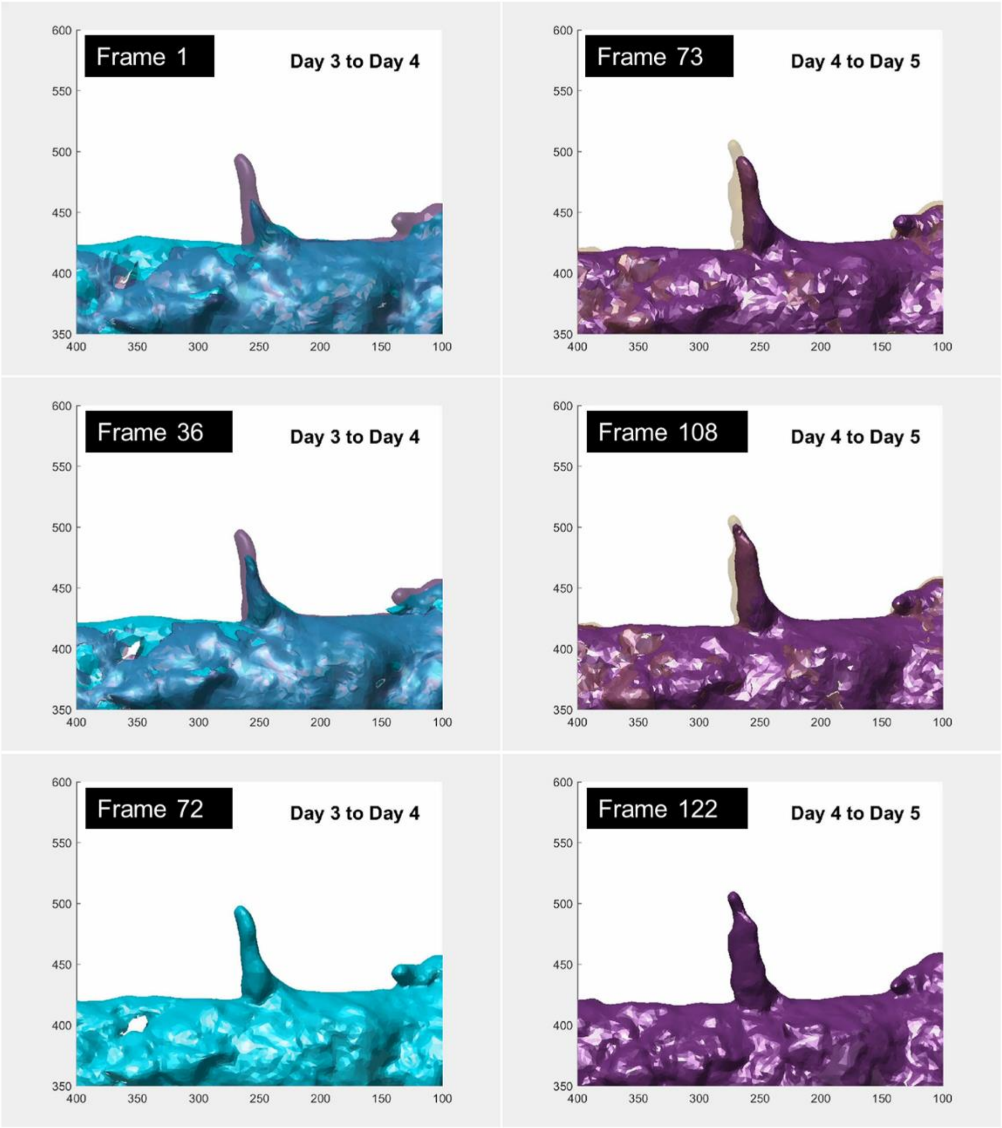


**Movie S1. Linear interpolation animation of an angiogenic sprout deformation.** Surface meshes from day 3 to day 5 of a representative angiogenic sprout was set and the animation frames were interpolated using the coordinates and the deformation vector between corresponding vertices in two continuous time points. Axes present x-axis (horizontal) and y-axis (vertical) of the mesh. Unit: μm.
